# Allosteric regulation of a molecular motor through *de novo* protein design

**DOI:** 10.1101/2023.10.17.562760

**Authors:** Jessica A. Cross, William M. Dawson, Shivam R. Shukla, Johannes F. Weijman, Judith Mantell, Mark P. Dodding, Derek N. Woolfson

**Affiliations:** School of Biochemistry, University of Bristol, University Walk, BS8 1TD, UK; School of Chemistry, University of Bristol, Cantock’s Close, Bristol BS8 1TS, UK; Bristol BioDesign Institute, University of Bristol, Cantock’s Close, Bristol BS8 1TS, UK

## Abstract

Many enzymes are allosterically regulated. Our ability to manipulate these structural changes is limited. Here we install an allosteric switch into the kinesin-1 microtubule motor *in vitro* and in cells. Kinesin-1 is a heterotetramer that accesses open active and closed auto-inhibited states. The equilibrium between these centres on a flexible elbow within a complex coiled-coil architecture. We target the elbow to engineer a closed state that can be opened with a *de novo* designed peptide. The alternative states are modelled computationally and confirmed by biophysical measurements and electron microscopy. In cells, peptide-driven activation increases kinesin transport, demonstrating a primary role for conformational switching in regulating motor activity. The designs are enabled by our understanding of ubiquitous coiled-coil structures, opening possibilities for controlling other protein activities.

**One Sentence Summary:** *De novo* peptide and protein design are used to engineer an allosteric switch into kinesin-1 motors *in vitro* and directly in cells.

## Introduction

Conformational switches are key regulators of protein function and activity. Specifically, allostery refers to a conformational change in a protein or protein complex brought about by binding an effector molecule to alter the protein’s function or activity. An ability to engineer allostery into natural and *de novo* designed proteins would have considerable impact in cell, chemical and synthetic biology. However, amongst other challenges, allosteric binding sites are often distant from the active sites of enzymes, which makes coupling the two, in successful designs, a complex endeavour.

The predominant conformation of a protein is determined by its lowest energy state for a given set of conditions. Conformational switching requires delicate control of the equilibrium between two or more states, with fine-tuned energetics to drive exchange in response to a stimulus. Whilst protein design is advancing rapidly (*1-3*), capturing such balances in *de novo* proteins is extremely challenging. Indeed, protein design has largely sought to maximise the free-energy difference between a single, desired, folded structure and the unfolded and alternative states. In this respect, the state of the art of protein design is impressive as defined single states are increasingly being delivered with high accuracy (*4-7*). However, many of these are hyperstable (*8-11*), likely less dynamic than natural protein structures, and thus limited for exploring conformational dynamics, alternate states, and allostery. Therefore, the design of more plastic and potentially switchable protein structures is a priority. Indeed, the field has sought to mimic protein conformational switches through multi-state design for some time (*3, 12, 13*); and there are examples of allosteric switches that have been engineered into natural proteins (*14*), or designed *de novo* (*15-19*). A new challenge is to apply developments from largely test-tube-based designs to manipulate complex natural proteins *in situ*.

Dynamics between different conformational states are particularly important in cytoskeletal motor proteins, which are involved in a range of cellular processes including intracellular transport, cell division, and cell migration. Motor activity and regulation are critical for healthy cell function and dysregulation can lead to disease (*20-22*). Therefore, strategies to manipulate motor-protein activity would progress fundamental cell biology and biomedical applications. A key aspect of motor regulation involves autoinhibitory interactions within the multi-domain/multi-subunit protein complexes.

For example, kinesin-1 is a ubiquitous microtubule motor involved in intracellular cargo transport within most eukaryotic cells, with a prominent role in axonal transport. It forms a heterotetramer comprising two motor-bearing heavy chains (KHCs) and two cargo-binding/regulatory light chains (KLCs) (*23*). When not transporting cargo, kinesin-1 enzymatic activity is inhibited by interactions between *C-*terminal regulatory motifs (isoleucine-alanine-lysine (IAK) motifs) and the *N-*terminal motor domains of the KHCs (Fig. 1A). Effectively, these interactions cross-link the motor domains, suppressing microtubule-dependent adenosine triphosphatase (ATPase) activity (*24-27*). Recently, we combined computational structure prediction with single-particle negative-stain electron microscopy (NS-EM) to visualise the complex coiled-coil architecture of the kinesin-1 autoinhibited state. This revealed a flexible hinge between two coiled-coil domains (CC2 and CC3) of KHC, ‘the elbow’. This region is critical for the complex to switch between an open state (Fig. 1B) and a folded-over ‘lambda’ particle, which brings the IAK motifs and motor domains together (Fig. 1A) (*28*). This is supported by experiments combining cross-linking and mass spectrometry (*29*).

**Fig. 1.**
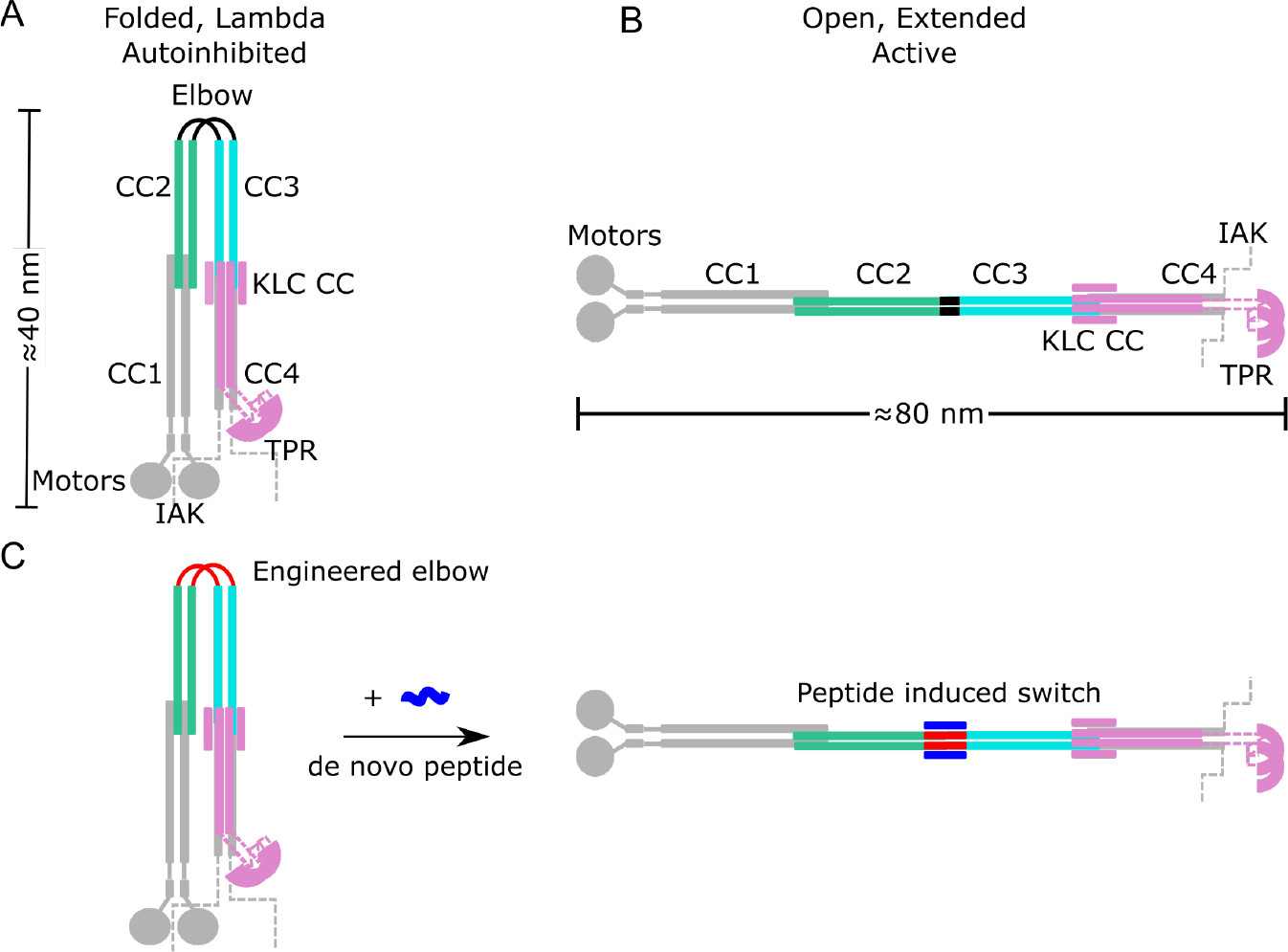
Architecture of the autoinhibited and active states of kinesin-1 and target design. **A, B**. Schematic illustrating the autoinhibited lambda particle (**A**) and open conformer (**B**) of heterotetrameric kinesin-1. Kinesin heavy chains (KHCs) are shown in grey with CC2 highlighted in teal and CC3 in cyan. Kinesin light chains (KLCs) are shown in purple. Globular domains and hinges are labelled and coiled-coil domains are numbered CC1 – 4. Grey dashed lines indicate the unstructured KHC *C*-terminal tails that contain the IAK regulatory motif. **C**. The aim of this work is to engineer a *de novo* peptide (red) into the elbow of a lambda state to construct a peptide-inducible (blue) conformational switch to the open state.

It is generally thought that the open form and the lambda particle are associated with the active and autoinhibited states of kinesin-1, respectively, but the extent of the conformational change and how it is regulated is unclear. We reasoned that this could be addressed through protein design to target the coiled coil-based switch. Coiled coils are attractive targets because the unprecedented understanding of them provides a strong basis for design (*30*). Indeed, the *de novo* design of static coiled coils with defined oligomeric states is considered largely solved (*1, 30*). Remaining challenges in coiled-coil design include making coiled coil-based switches (*31-36*), and using our understanding to intervene in biological processes directly *in situ*.

Here we address these questions in kinesin cell biology and challenges in protein design to deliver *de novo* protein-peptide interactions that control the kinesin-1 motor in cells. To do this, we apply protein design principles to re-engineer the motor-activation switch and render it activatable by a *de novo* designed cell-penetrating peptide (Fig. 1C). This establishes a clear role for a large-scale conformational change pivoting on the elbow to regulate motor activity; and demonstrates the potential to manipulate coiled-coil switches in biology to understand and hijack protein function.

## Results

### Kinesin-1 conformations can be controlled by protein engineering and design

For recombinantly expressed and purified Kinesin^WT^ (composed of His-Kif5C (KHC) and KLC1 isoform A (KLC)), the lambda particle and open state (Fig. 1A-B) are apparent as two peaks in size-exclusion chromatography (SEC) (Fig. 2A), and as two conformations in negative stain electron microscopy (NS-EM) (*28*). Recently, using knowledge of coiled-coil folding and assembly (*30, 37*), we have engineered an 18-residue elbow deletion, Kinesin^Delta Elbow^, to promote helical readthrough from CC2 to CC3 and favour the open state (Fig. 2B), which is then observed as a single peak in SEC and can be visualised by NS-EM (*28*). As a platform to build a switch for this new study, we sought other engineered variants that stabilise the two states of the Kinesin^WT^ complex using a combination of rational protein engineering and design, modelling with AlphaFold2 (*38, 39*), and SEC as the initial experimental reporter.

**Fig. 2.**
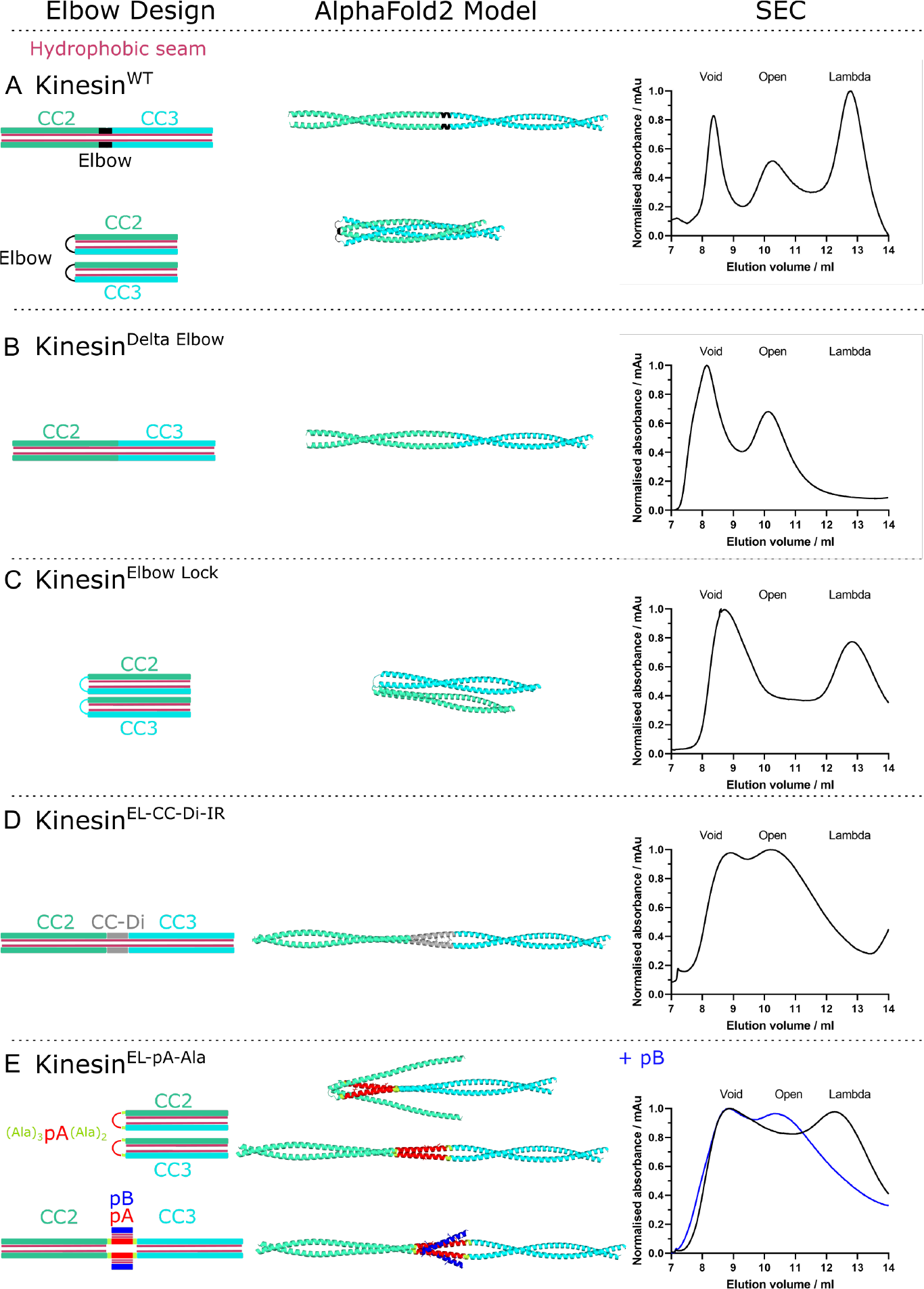
Engineering an allosteric switch into kinesin-1 that is activated by a *de novo* designed peptide. From left to right: cartoon illustrations of each elbow design; AlphaFold2 models of KHC for the CC2-linker-CC3 region; and size-exclusion chromatography (SEC) elution profiles of purified heterotetrameric KHC-KLC complexes without (black) and with (blue) peptide pB. Colour scheme for the structural cartoons: KHC CC2 teal and CC3 cyan; hydrophobic seams/cores, pink; CC-Di, grey; peptide pA, red; flanks, yellow; and peptide pB, blue. **A**. Wild-type kinesin-1 (Kinesin-1^WT^) exists in an equilibrium between the open and lambda states. **B**. The Delta Elbow variant of kinesin-1 (Kinesin-1^Delta Elbow^), with an 18-residue deletion in the elbow region, favours the open state. **C**. The Elbow Lock variant (Kinesin-1^Elbow Lock^), which has a five-residue deletion in the elbow region, favours the lambda state. **D**. Introduction of a 28-residue *de novo* designed peptide CC-Di into the deleted five-residue elbow of kinesin-1 (Kinesin-1^EL-CC-Di-IR^) favours the open state. **E**. Designed switchable variant Kinesin^EL-pA-Ala^ without (top) and with (bottom) peptide pB. Insertion of pA with a shifted coiled-coil register relative to CC2 and CC3 using helix-favouring (Ala)_3_ and (Ala)_2_ flanks, respectively, allows flexibility in the elbow without pB favouring the lambda particle, but induced coiled-coil formation with pB activates the switch to the open state. Alphafold models predict both an open and alternative compact state without peptide pB but all open models with pB.

First, we tested if AlphaFold2 (*38, 39*) could model KHC variants reliably. AlphaFold2 predictions of the homodimer for the WT CC2-elbow-CC3 region appeared to capture the two-state equilibrium. Here, the standard five AlphaFold2 models included a fully extended state in which CC2, the elbow, and CC3 form a single helix that dimerises with an unbroken hydrophobic core, which we interpret as analogous to the open state; and four predictions of a state in which the elbow forms a loop allowing CC2 and CC3 dimers to fold back on each other (Fig. 2A and S1). The latter closely resembles our AlphaFold2 model for the full coiled-coil architecture of the heterotetrameric lambda particle (*28*). Moreover, AlphaFold2 models for the corresponding region of the Kinesin^Delta Elbow^ variant of KHC were all fully extended coiled coils (open state) consistent with our previous experiments (Fig. 2B and S2) (*28*).

To address our first objective of engineering a variant that stabilises the folded lambda particle, we used Socket2 (*40*) to analyse the interhelical, coiled-coil, knobs-into-holes interactions predicted around the elbow of the full AlphaFold2 model for the lambda particle (*28*). This indicated that a five-residue deletion from the loop, Kinesin^Elbow Lock^, might stabilise the folded-over conformation by disrupting the coiled-coil, heptad, sequence repeats and reducing conformational flexibility (Table S1). Indeed, this modified KHC sequence returned only AlphaFold2 models compatible with the lambda particle and no extended (open) forms (Fig. 2C and S3). Moreover, when expressed and co-purified with KLC, Kinesin^Elbow Lock^ gave a complex that eluted as a single peak for the lambda particle, indicating that the equilibrium had indeed been completely shifted to the compact state (Fig. 2C).

Next, we asked if these key five-residues in the elbow could be replaced to design a switch back to the open conformation. Our strategy was to insert a stable coiled coil into the target (*18*). Specifically, we used the 28-residue, homodimeric, *de novo* coiled coil, CC-Di (*41*). We inserted this sequence into the deleted elbow region to generate Kinesin-1^EL-CC-Di-IR^, with a contiguous (in-register, IR) hydrophobic core bridging the CC2, inserted coiled coil, and CC3 domains (Fig. 2D and Table S1). Consistent with this, AlphaFold2 models of Kinesin-1^EL-CC-Di-IR^ were solely for the extended form. Furthermore, experimentally, this complex eluted exclusively as the open state, (Fig. 2D and S4).

### A *de novo* designed peptide induces a switch between the lambda and open states

With engineered lambda and open conformations of kinesin-1 in hand, we targeted the design of a switch between these that could be triggered by a synthetic *de novo* designed peptide. The logic was to swap CC-Di in Kinesin-1^EL-CC-Di-IR^ for a heterodimeric coiled coil, the formation of which would drive coiled-coil extension leading to the open state and activation of the motor, (Fig. 1C). We chose our heterodimeric antiparallel coiled coil, apCC-Di-AB (*42*), which has two components referred to here as pA and pB (Table S1). Individually, pA and pB are unfolded, but when combined they assemble into a stable coiled-coil heterodimer (*42*). We sought to design a construct that introduced pA into the five-residue elbow deletion, retaining the flexibility between CC2 and CC3 required to form the lambda particle, and keeping pA accessible for subsequent binding by pB.

Initially, we introduced pA in the same heptad register as CC2 and CC3 (Kinesin-1^EL-pA-IR^), analogous to Kinesin-1^EL-CC-Di-IR^ (Table S1). Although pA is unfolded in isolation, when templated between CC2 and CC3 in this way, the sequence was predicted by AlphaFold2 to form an extended coiled coil (Fig. S5). This was confirmed by SEC with the complex mostly in the open state (Fig. S6A). Binding of pB to pA was not modelled well by AlphaFold2 (Fig. S6A and S7), but was confirmed experimentally by measuring the co-elution of fluorescently labelled peptide with the KHC-KLC heterotetrameric complex in SEC (Fig. S8). Binding did not yield an allosteric switch, which we attribute to the largely open population of this engineered kinesin in the absence of pB. In an attempt to address this, we introduced flexibility either side of the pA insert to prevent helical readthrough and templating by the flanking CC2 and CC3 domains. First, we tested triple-glycine linkers either side of pA (Kinesin-1^EL-pA-Gly^; Table S1, Fig. S6B and S9). SEC confirmed a compact state for the resulting complex (Fig. S6B). AlphaFold2 predicted that Kinesin-1^EL-pA-Gly^ could bind pB as designed but still in a folded conformation, (Fig. S10). Binding of the peptide was confirmed by the fluorescence-based assay (Fig. S11). However, although this second design iteration favoured compact states of the kinesin complex, again, it did not enable the targeted switch.

The lessons taken from these designs were that a break in coiled-coil/helical readthrough is required, but without excessive structural flexibility either side of the insert. Therefore, we placed helix-favouring alanine residues strategically either side of pA (three alanine residues preceding it and two after it) to shift its hydrophobic face by ≈180° relative to those of the kinesin-1 CC2 and CC3 helices (Table S1, Fig. 2E). Thus, in this design, Kinesin-1^EL-pA-Ala^, the hydrophobic *a* and *d* sites of the *a – g* heptad repeats of pA should fall at solvent-exposed *f* and *b* positions of the kinesin-1 coiled-coil registers. This should not template folding of the pA insert. Without peptide pB, AlphaFold2 did not model the KHC dimer well and both compact (1/5) and open (4/5) models were returned (Fig. S12). Encouragingly, however, exclusively open models were predicted for the pB-bound Kinesin-1^EL-pA-Ala^ dimer (Fig. 2E and S13). Fully consistent with our design rationale, SEC revealed that the switch could be fully realised for this design, (Fig. 2E and S14): without pB, the major complex eluted at the same volume as the lambda particle; and with pB the elution profile shifted dramatically to the open state.

### The open and lambda states can be visualised directly *in vitro*

To examine the conformational state of kinesin-1 complexes in solution, we collected SAXS data in-line with SEC (SEC-SAXS) (*43*). From the scattering profiles (Fig. S15), we derived the radius of gyration (*R*_g_) and maximum diameter for each particle (*D*_max_) and generated bead models using DAMMIN (*44*). Analysis of Kratky plots indicated flexibility in all the kinesin complexes (Fig. S16). Notably, two different species were apparent for the Kinesin^WT^ heterotetramer, with *D*_max_ values of 45 nm and 65 nm, respectively (Fig. S17, Table S2). These are indicative of the open state and lambda particle, respectively (*28*). Consistent with the predictions and SEC profiles above, Kinesin^Delta Elbow^ and Kinesin^EL-CC-Di-IR^ gave a single extended conformation of *D*_max_ ≈ 70 nm, like the open state. In contrast, Kinesin^Elbow Lock^ and Kinesin^EL-pA-Ala^ showed single conformations of *D*_max_ ≈ 40 nm, indicating the lambda conformation (Fig. 3A-D, Table S2). Moreover, addition of pB to Kinesin^EL-pA-Ala^ caused a shift in *D*_max_ from ≈ 40 nm to ≈ 70 nm (Fig. 3E, Table S2). This provides direct biophysical evidence that binding of pB induces a conformational change in Kinesin^EL-pA-Ala^ from a closed to an open conformation.

**Fig. 3.**
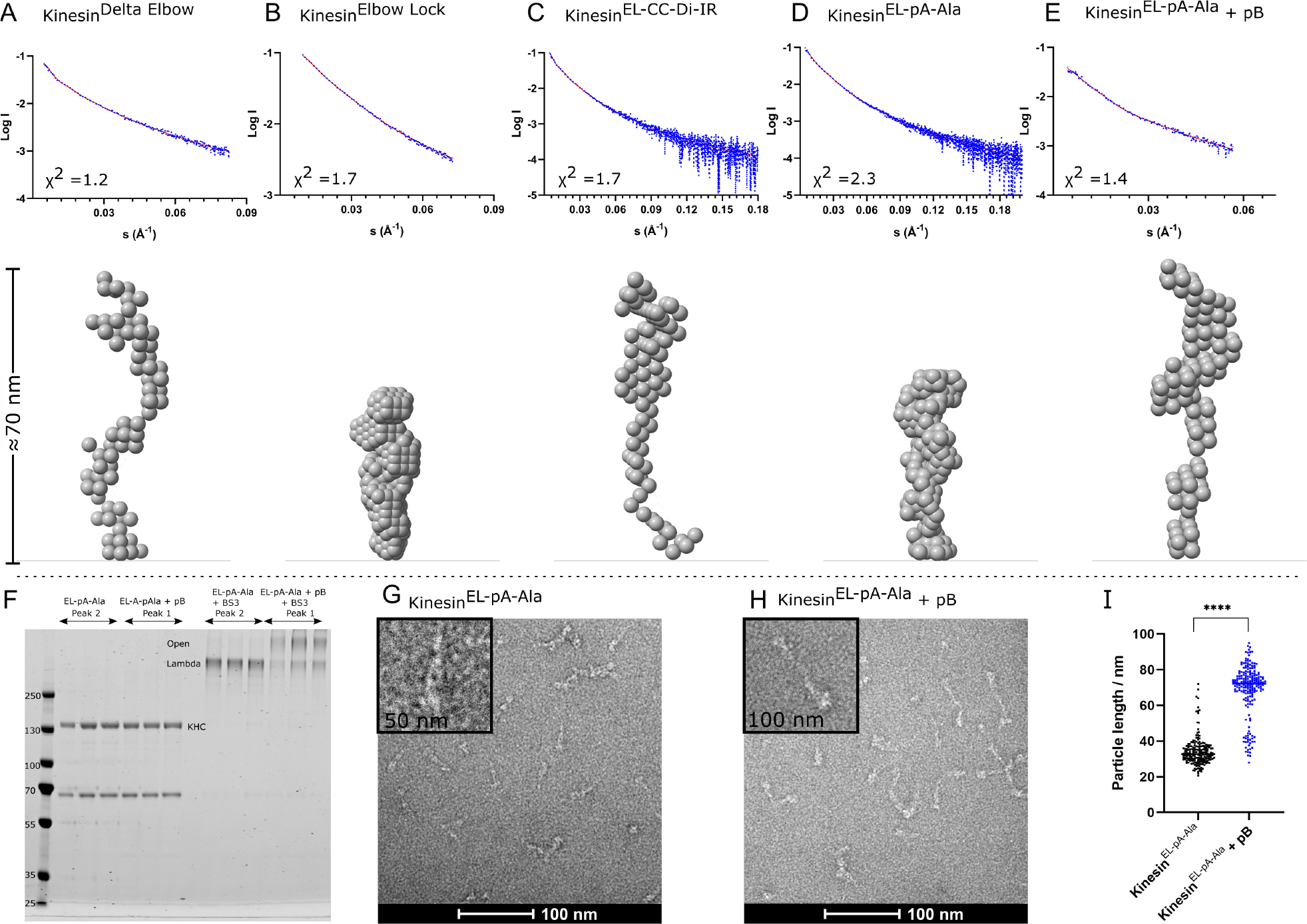
*In vitro* visualisation of the lambda and open conformational states of kinesin-1 elbow variants. **A-E**. SEC-SAXS in-solution shape determination. Top: Comparison of trimmed experimental scattering profiles (blue) with theoretical scattering profiles (red) generated from *ab initio* models using DAMMIN. The quality of fit is assessed using the χ^2^ statistic. Bottom: Representative models obtained from DAMMIN illustrating the shapes of the kinesin-1 variants. **A**. Kinesin-1^Delta Elbow^ is in the open state *D*_max_ = 70 nm χ^2^ = 1.7, **B**. Kinesin-1^Elbow Lock^ is in the lambda state *D*_max_ = 38.3 nm χ^2^ = 1.2, **C**. Kinesin-1^EL-CC-Di-IR^ is in the open state *D*_max_ = 69.7 nm χ^2^ = 1.7, **D**. Kinesin-1^EL-pA-Ala^ is in the lambda state *D*_max_ = 40 nm χ^2^ = 2.3, **E**. Kinesin-1^EL-pA-Ala^ + pB is in the open state *D*_max_ = 69 nm χ^2^ = 1.4. **F-I**. Negative-stain electron microscopy visualises the peptide-induced conformational switch of kinesin-1. **F**. Coomassie-stained SDS-PAGE gel showing complexes containing Kinesin-1^EL-pA-Ala^ purified by SEC. Complexes eluted in two distinct peaks after the void for kinesin-1 without (EL-pA-Ala, peak 2) and with pB (EL-pA-Ala + pB, peak 1) sampled with 0.25 ml fractions through these peaks. Lanes marked with +BS3 show the mobility of duplicate samples after cross-linking. Cross-linked complexes formed in the presence of pB migrate slower through the gel indicating larger (open) particles. **G, G** Representative T12 electron micrographs showing negative-stained, cross-linked samples from EL-pA-Ala (**G**) and EL-pA-Ala + pB (**H**). Scale bars, 100 nm. Zoom box size is 50 nm (**G**) and 100 nm (**H**). **I**. Quantification of particle length from panels **G** and **H** for 100 particles from each sample. Addition of pB (blue) results in an increase in the length of particles consistent with a switch from the lambda particle to the open state.

As an orthologous approach to confirm the peptide-induced switching of Kinesin-1^EL-pA-Ala^, we trapped and visualised its states with and without pB. The protein complexes were isolated as the two major peaks from SEC and treated with the cross-linker bis(sulfosuccinimidyl)suberate (BS3) (*28*). In SDS–polyacrylamide gel electrophoresis (PAGE), the complex alone gave a faster-migrating species compared to that with pB, (Fig. 3F). These data indicate further that Kinesin-1^EL-pA-Ala^ alone is more compact consistent with the lambda particle, whilst the complex formed with pB is more elongated. This was confirmed by NS-EM, which revealed that the major species for Kinesin-1^EL-pA-Ala^ alone resembled our previously described Kinesin-1^WT^ lambda particle (*28*); namely, an ≈ 40 nm-long V-shaped object with wide and tapered ends (Fig. 3G, I). In contrast, with pB, Kinesin-1^EL-pA-Ala^ gave predominantly long or slightly bent and thinner particles ≈ 70 – 80 nm in length, similar to the open conformers of Kinesin-1^WT^ and Kinesin-1^Delta Elbow^ (Fig. 3H-I). Thus, the engineered Kinesin-1^EL-pA-Ala^ complex can access both the closed (lambda) and open states and addition of the complementary *de novo* designed peptide, pB, activates the switch to the latter.

### The allosteric switch can be triggered in cells with pB applied exogenously

Finally, we tested if the peptide-induced conformational switch could allosterically activate kinesin-1 in cells. HeLa cells were transiently transfected with two genes to express kinesin-1 heterotetramers. For visualisation, KLC was conjugated to a green fluorescent protein (GFP-KLC2) and KHC to a haemagglutinin tag (HA-Kif5C). The KHC gene was mutated to incorporate the elbow variants and then imaged to examine kinesin-1 localisation by fluorescence light microscopy. Cells expressing Kinesin-1^WT^ and Kinesin-1^Elbow Lock^, which favour the lambda-particle conformation, displayed diffuse cytosolic localisation of both KHC and KLC characteristic of autoinhibited kinesin. By contrast, Kinesin-1^EL-CC-Di-IR^, which favours the open state, gave a different phenotype of extended cells with peripheral accumulations of kinesin-1 (Fig. 4A). This indicates that this variant is a constitutively active kinesin motor that moves along microtubules to the cell periphery. pB is also a cell-penetrating peptide that enters directly into the cytoplasm and nuclei of HeLa cells (*42*). In cells expressing

**Fig. 4.**
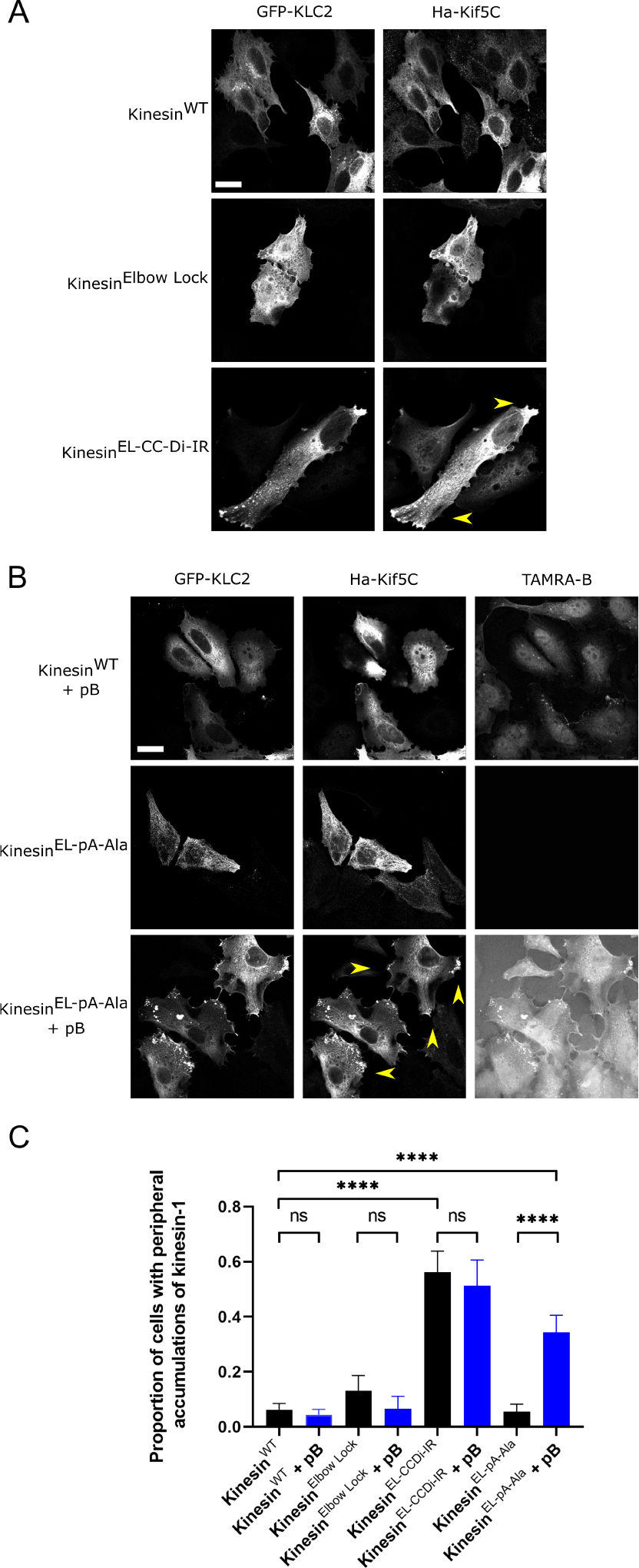
Distribution of kinesin-1 complexes in HeLa cells. **A**. HeLa cells transfected with GFP-KLC2 and Ha-Kif5C (Kinesin-1^WT^, Kinesin-1^Elbow Lock^ or Kinesin-1^EL-CC-Di-IR^). Kinesin-1^WT^ and Kinesin-1^Elbow Lock^ complexes show diffuse cytosolic localisation indicative of autoinhibition. Kinesin-1^EL-CC-Di-IR^ displays extended cell morphology and peripheral accumulations of kinesin-1 at cell vertices. Scale bar, 20 μm. **B**. HeLa cells transfected with GFP-KLC2 and Ha-Kif5C (Kinesin-1^WT^, Kinesin-1^EL-pA-Ala^) with/without treatment with 2 μM TAMRA-labelled pB for 1 h. pB is cell penetrating as shown by TAMRA fluorescence in the cytoplasm and nucleus of Kinesin-1^WT^ transfected cells. pB binds to and colocalises with Kinesin-1^EL-pA-Ala^ complexes in peripheral accumulations at cell vertices. Scale bar, 20 μm. **C**. Quantification of proportion of cells with peripheral accumulations of kinesin-1 in panels **A** and **B** in a minimum of 45 cells pooled from three independent experiments. Additional controls of Kinesin-1^Elbow Lock^ and Kinesin-1^EL-CC-Di-IR^ with pB show no change on addition of peptide. Data are presented as mean ± S.E.M. and an unpaired t-test is used for statistical analysis. ***P < 0.0001.

Kinesin-1^EL-pA-Ala^, adding fluorescently labelled pB to the media resulted in its co-localisation with KLCs and KHCs and caused a dramatic redistribution of kinesin-1 complexes to peripheral accumulations (Fig. 4B). In control experiments without pB, Kinesin-1^EL-pA-Ala^ resembled the diffuse distribution of Kinesin-1^WT^ and Kinesin-1^Elbow Lock^. Together, these experiments show that cell-penetrating pB can target the engineered Kinesin-1^EL-pA-Ala^ to effect acute allosteric activation in living cells (Fig. 4B-C).

## Discussion

In summary, we have applied rational protein design principles to engineer a peptide-induced allosteric switch within a molecular motor that can be triggered *in vitro* and in cells using an exogenous reagent. In doing so, we show how a fundamental understanding of sequence-to-structure relationships of coiled-coil domains can enable the design of reagents to target them in complex biological machines.

This design-led approach also provides important insight on the mechanism of kinesin-1 autoregulation and, going forwards, our ability to target it. The capacity to stabilise or to relieve autoinhibition *via* subtle manipulations of the elbow region indicates that this hinge is a key and finely tuned regulatory switch of the kinesin-1 motor complex. Indeed, introduction of a *de novo* designed coiled-coil peptide with nM affinity (*42*) is sufficient to switch the conformation from the folded lambda particle to an open state and promote activity. From this, we posit that the sum of all other interactions involved in autoinhibition must be less than the free energy change associated with the designed binding event. Therefore, it seems that the primary function of the motor-IAK autoinhibitory interactions is not to drive closure to the lambda state, but rather to supress enzymatic activity and modulate cargo-binding properties within this state. This notion is consistent with recent observations that mutation of the IAK motif is not sufficient to open and activate the complex (*29, 45*).

In nature, kinesin-1 motors are activated through binding of a diverse array of cargo *via* adaptor proteins (*26, 46*), but the underlying mechanisms remain unclear. A key inference from our study is that the primary function of many coiled-coil-binding cargo-adaptor proteins and microtubule-associated proteins (such as MAP7) is to perturb the finely balanced equilibrium between the lambda and open state (*26, 47, 48*). Our findings indicate that the autoinhibitory mechanism based around this coiled-coil switch is poised to respond to both endogenous and exogenous inputs. Therefore, it may be amenable to designing new reagents that target the endogenous protein to boost or supress kinesin-1 mediated intracellular transport where it is implicated in disease.

In addition, the constructs that we have derived will offer valuable tools for future studies into both the structure and function of kinesin-1 by stabilising either the autoinhibited or active state without removing key cargo recognition and regulatory components of the protein complex. Our system offers an acute means for in-cell activation of kinesin-1 without perturbing key cargo binding and enzymatic activities.

A long-standing question is the extent to which the kinesin-1 complex opens to engage in cargo transport. Although most models assume that the complex fully extends, others suggest that a more-compact form is responsible for cargo transport (*49, 50*). Our finding that acute peptide-driven extension is sufficient to activate the complex favours the extension model. Our ability to make and test model-based predictions for successful protein design, indicates that the elbow sequence is highly evolved to adopt two distinct states – a flexible hinge and a rigid read-through of helical, heptad repeats from CC2 to CC3. Our findings also suggest limited flexibility in the rest of the coiled-coil scaffold. We propose that rigid read-through (modulated by cargo adaptor binding) is most likely to represent the functional active complex.

The kinesin-1 motor system offered an attractive target to test our engineering and design approach to effect an allosteric switch in a natural enzyme. We believe that this approach may be developed in various ways. Firstly, the principles that we have devised should be applicable to other proteins whose conformational state is regulated by hinging coiled coils that can switch between loop and folded conformations (*18, 51-55*). Secondly, reagents of the type we present could be developed to target natural coiled-coil domains more generally. This work demonstrates the importance of balancing interaction affinities within protein complexes and with their effectors. Potentially, our approach ushers in a new era of quantitative chemical biology. If realised, this would open the door to larger scale efforts to target these domains, which comprise up to 15% of the human proteome (*56, 57*).

## Acknowledgements

J.A.C. is supported by an EPSRC-funded Doctoral Prize Fellowship. This work was supported by a BBSRC grant to M.P.D. and D.N.W (BB/W005581/1). M.P.D. acknowledges support from a Lister Institute of Preventative Medicine Fellowship. W.M.D. and D.N.W. were funded by a European Research Council Advanced Grant (340764) and a subsequent European Research Council Proof of Concept Grant (787173). D.N.W. was also supported by the BrisSynBio, a BBSRC/Engineering and Physical Sciences Research Council-funded Synthetic Biology Research Centre [BB/L01386X/1]. We thank the University of Bristol School of Chemistry Mass Spectrometry Facility for access to the EPSRC-funded Bruker Ultraflex MALDI-TOF instrument (EP/K03927X/1), the BBSRC-funded BrisSynBio centre for access to peptide synthesis and the plate reader (BB/L01386X/1), and the Wolfson Bioimaging Facility for access to the confocal and electron microscopes and their assistance in this work. We acknowledge financial support and allocation of beamtime at Diamond Light Source and we thank the beamline staff at B21, in particular Dr Nathan Cowieson and Dr Nikul Khunti, for assistance.

## Author contributions

J.A.C, W.M.D, M.P.D and D.N.W conceived and developed the project. J.A.C, W.M.D, M.P.D and D.N.W designed the proteins and conducted *in silico* analysis. J.A.C, S.R.S, J.F.W and J.M. prepared and the proteins and conducted biophysical analysis. J.A.C. conducted mammalian cell experiments. J.A.C, W.M.D, S.R.S, M.P.D and D.N.W conducted formal analysis. J.A.C., M.P.D and D.N.W. wrote the manuscript with contributions from all authors.

