## supplementary materials for "Allosteric regulation of a molecular motor through *de novo* protein design"

[3]Bristol BioDesign Institute, University of Bristol, Tyndall Avenue, Bristol BS8 1TQ, UK

**Supplementary Materials**
Materials and Methods
Tables S1 to S3
Figs. S1 to S17
References (*1-11*)

### Materials and Methods

#### AlphaFold2 predictions

Predictions were made using the Google Colab Alphafold Notebook with a Colab Pro+ subscription (*1, 2*). The algorithm was asked to model a 1:1 homodimer composed of rat KIF5C (NP_001101200.1) residues I590 to L772 or a 2:2 heterotetramer of these sequences with peptide pB. These boundaries were established by combining insight from Marcoil (*3, 4*) predictions and Socket-2 (*5*) analysis of a larger model described previously (*6*). Default parameters were used, and the amber relaxation step was enabled. The five output models were downloaded in pdb format, aligned and prepared for presentation using PyMOL. Model statistics are as presented by the software.

#### Plasmids used in this study

Previously, rat KIF5C (corresponding to residues 2 to 955), was amplified by polymerase chain reaction (PCR) from an HA-tagged expression construct and ligated into the His-3C cleavage pMW bacterial expression vector and mouse KLC1 was amplified by PCR and ligated into pET28a using Nde I/Xho I sites with the N-terminal His-thrombin cleavage tag removed by site-directed mutagenesis (*6, 7*). Clonal genes containing fragments of the elbow designs described in this study were synthesised by Twist Bioscience and cloned into pMW-His-rat Kif5C using BrsG I / EcoR 1 sites. These constructs were subcloned into a CB6 mammalian expression vector using Not I/EcoR I sites.

### Protein expression and purification

Heterotetrametric kinesin-1 was expressed in BL21(DE3) cells using a two-plasmid system. BL21(DE3) cells, transformed with both KIF5C and KLC1 expression plasmids, were used to inoculate 1 litre of LB cultures supplemented with both ampicillin and kanamycin. Cells were grown with shaking at 37°C until optical density reached 0.8 before the cultures were cooled to 18°C and protein expression was induced with 0.3 µM isopropyl-β-d-thiogalactopyranoside. Following shaking incubation at 18°C overnight, cells were harvested by centrifugation at 6000 *g* at 4°C for 15 min and resuspended (10 ml per 1 litre of original culture) in 20 mM Hepes (pH 7.4), 300 mM NaCl, and 40 mM imidazole before being stored at −20°C. Frozen pellets were thawed and diluted in 25 ml of buffer consisting of 40 mM Hepes (pH 7.4), 500 mM NaCl, 40 mM imidazole, 5% (v/v) glycerol, and 5 mM β-mercaptoethanol with a Roche complete protease inhibitor tablet added. Bacteria were lysed by sonication, 0.5 s on and 10 s off, at 70% amplitude for 7 min and 30 s in an ice bath. Lysate was clarified by centrifugation at 35,000 *g* on a JA-20 rotor for 40 min at 4°C. Clarified lysate was filtered (0.45 µm) before being loaded onto a His-Trap (Sigma-Aldrich) column. The column was washed in buffer and eluted with a gradient of 40 to 500 mM imidazole. Eluted protein was concentrated by ultrafiltration in a 10,000-Da molecular weight cutoff filter (Cytiva), before snap-freezing in liquid nitrogen.

### Size exclusion chromatography

Proteins were thawed and incubated for 1h at 4°C with agitation in the presence of peptide pB or a vehicle control (1 µl $H_2O$). Proteins were further purified by SEC using a Superose6 10/300 column (Cytiva), in 20 mM Hepes (pH 7.4), 150 mM NaCl, 1 mM $MgCl_2$, 0.1 mM adenosine 5′-diphosphate, and 0.5 mM tris(2-carboxyethyl)phosphine (Sigma-Aldrich) at a flow rate of 0.5 ml/min. Elution of protein was monitored by following absorbance at 280 nm and individual fractions were run on a Coomassie stained gel to confirm size and purity. Elution of peptide pB was monitored by measuring fluorescence at 555 nm of individual fractions in a plate reader. Measurements were repeated a minimum of three times and one representative trace is shown for clarity.

### Peptide synthesis

Peptide pB was prepared by standard Fmoc solid-phase peptide synthesis on a 0.1 mM scale using CEM Liberty Blue automated peptide synthesis apparatus with inline UV monitoring. Activation was achieved with DIC/Oxyma. Fmoc deprotection was performed with 20% v/v morpholine/DMF. Double couplings were used for β-branched residues and the subsequent amino acid. Synthesis was from C to N terminus as the C-terminal amide on Rink amide resin and labelling achieved by addition of TAMRA (0.1 mM, 2 eq.), HATU (0.095 mM, 1.9 eq.) and DIPEA (0.225 mM, 4.5 eq.) in DMF (3 mL) to DMF washed peptide resin (0.05 mM) with agitation for 3 hours. Resin was washed with 20% piperidine in DMF (5 mL) for 2 x 30 minutes to remove any excess dye. All manipulations were carried out under foil to exclude light. Peptide pB was cleaved from the solid support by addition of TFA (9.5 mL), TIPS (0.25 mL) and water (0.25 mL) for 3 hours with shaking at rt. The cleavage solution was reduced to ≈1 mL under a flow of nitrogen. Crude peptide was precipitated upon addition of ice-cold diethyl ether (40 mL) and recovered via centrifugation. The resulting precipitant was dissolved in 1:1 acetonitrile and water (≈15 mL) and lyophilised to yield crude peptide as a solid.

**Peptide purification**

Peptide pB was purified by reverse phase HPLC on a Phenomenex Luna C18 stationary phase column (150 x 10 mm, 5 µM particle size, 100 A pore size) using a preparative JASCO HPLC system. A linear gradient of 20–80% acetonitrile and water (with 0.1% TFA) was applied over 30 minutes. Chromatograms were monitored at wavelengths of 220 and 280 nm. The peptide was confirmed using MALDI-TOF mass spectrometry using a Bruker ultrafleXtreme II instrument in reflector mode. Peptide was spotted on a ground-steel target plate using a-cyano-4-hydroxycinnamic acid (CHCA) as the matrix. Masses were measured to 0.1% accuracy. Peptide purity was determined using a JASCO analytical HPLC system, fitted with a reverse-phase Kinetex® C18 analytical column (100 x 4.6 mm, 5 µm particle size, 100 Å pore size). Fractions containing pure peptide were pooled and lyophilised. Peptide was dissolved in buffer and concentration determined by UV–Vis at 280 nm on a ThermoScientific Nanodrop 2000 spectrophotometer by measurement of UV absorbance at 555 nm ($\varepsilon_{555}$(TAMRA) = 85000 $mol^{-1}cm^{-1}$).

**Small-angle X-ray scattering**

Small-angle X-ray scattering (SAXS) data were collected at Diamond Light Source Ltd. Synchrotron (Didcot, Oxfordshire, UK) on the B21 beamline, with an HPLC system upstream (Table S3) (*8*). As before, a superose6 10/300 column (Cytiva), was equilibrated in 20 mM Hepes (pH 7.4), 150 mM NaCl, 1 mM $MgCl_2$, 0.1 mM adenosine 5′-diphosphate, and 0.5 mM tris(2-carboxyethyl)phosphine (Sigma-Aldrich). Purified protein (95 ml of Kinesin-$1^{WT}$ 73 µM, Kinesin-$1^{Delta\ Elbow}$ 31 µM, Kinesin-$1^{Elbow\ Lock}$ 41 µM, Kinesin-$1^{EL\text{-}CC\text{-}Di\text{-}IR}$ 23 µM or Kinesin-$1^{EL\text{-}pA\text{-}Ala}$ 26 µM or Kinesin-$1^{EL\text{-}pA\text{-}Ala + pB}$ 20 µM)) was injected onto the pre-equilibrated column. The flow rate of the column was maintained at 0.5 mL/minute with eluted samples exposed to X-rays for 3 second exposure time Data were analysed in CHROMIXS (*9*). Further data processing was performed in PRIMUSQT (*10*). Ab-initio modelling was performed using DAMMIN (*11*).

**Negative-stain electron microscopy**

For negative stain electron microscopy, freshly eluted proteins from size exclusion were cross-linked with 0.6 mM BS3 (Thermo Fisher Scientific) for 30 min at room temperature. Cross-linked proteins were diluted to 0.003 mg/ml in size exclusion buffer, and 5 µl was pipetted onto a freshly glow-discharged grid (300-mesh copper with formvar/carbon support, TAAB) and incubated at room temperature for 1 min. Grids were manually blotted, washed in 3% uranyl acetate (UA) and stained in 20 µl of 3% UA for 20 seconds ,followed by a final wash in 3% UA, with the excess blotted away. The samples were air dried. Micrographs of grids were acquired on a FEI 120-kV BioTwin equipped with an FEI Ceta 4k x 4k charge-coupled device camera at ×49,000 magnification corresponding to a pixel size of 2.04 Å/pixel. Particle length was measured using ImageJ from several micrographs for each complex.

### Cell culture

HeLa cells were maintained in high glucose Dulbecco's Modified Eagle's Medium (Gibco Invitrogen) with 10% (v/v) foetal calf serum (Sigma-Aldrich) and 5% penicillin/streptomycin (PAA) (herein referred to as DMEM) at 37 °C and 5% $CO_2$. For transfection, cells were seeded in 6-well plates on fibronectin coated 13 mm coverslips at a density of 1 x $10^5$ cells per well and incubated at 37 °C and 5% $CO_2$ for 16 h prior to transfection. Cells were transfected with 0.4 µg DNA using Effectene transfection reagent according to the manufacturer's instructions (Qiagen). After transfection cells were incubated at 37 °C, 5% $CO_2$ for 16 hours. Peptide treatments were in 1 ml DMEM, 37 °C and 5% for 1 h with 2 µM TAMRA-pB or vehicle control (1 µl $H_2O$). Cells were fixed by addition of 4% paraformaldehyde in PBS at room temperature for 10 minutes (2 ml per well for 6-well plate) and washed 3 x with PBS. Confocal images were collected using a Leica SP5II system with a 63× objective running Leica LAS X and are presented as maximum intensity projections. Fig.s were assembled using ImageJ in conjunction with Inkscape. Image analysis was conducted in ImageJ.

**Table S1: Coiled coil register assignment of elbow designs used in this study**

| **Kif5C CC2 674→**<br>***abcdefg abcd***<br>EKMHEVS FQDK | **Elbow** | **CC3 → 698**<br>***cd abcdefg***<br>TR LQDAEEV |
|---|---|---|
| Kinesin$^{WT}$ | EKEHL | |
| Kinesin$^{Elbow\ Lock}$ | - | |
| Register$^{CC2/CC3/AB}$ | ***ef gabcdef gabcdef gabcdef gab*** | |
| Kinesin$^{EL-CC-Di-IR}$ | KQ EIAAIKK ENAALKW EIAALKQ EIA | |
| Kinesin$^{EL-A-IR}$ | EQ ELAALDQ EIAAAEQ ELAALDW QIQ | |
| Kinesin$^{EL-A-Gly}$ | *GGG* QLEQ ELAALDQ EIAAAEQ ELAALDW QIQ *GGG* | |
| Register$^{CC2/CC3}$ | ***efgabcd efgabcd efgabcd efgabcd efgab*** | |
| Kinesin$^{EL-A-Ala}$<br>Register$^{AB}$ | *AAA*QLEQ ELAALDQ EIAAAEQ ELAALDW QIQ*AA*<br>***cdef gabcdef gabcdef gabcdef gab*** | |
| Peptide-B | *G* QLKQ RRAALKQ RIAALKQ RRAALKW QIQ *G* | |

**Table S2. Size distribution parameters $D_{max}$ and $R_g$ values obtained from distance distribution ($P(r)$) analysis**

| **Elbow Design** | **$D_{max}$ (nm)** | **$R_g$ (nm)** |
|---|---|---|
| Kinesin$^{WT:\ Lamda}$ | 45.3 | 13.7 |
| Kinesin$^{WT:\ Open}$ | 64.7 | 16.7 |
| Kinesin$^{Delta\ Elbow}$ | 70 | 22.2 |
| Kinesin$^{Elbow\ Lock}$ | 38.3 | 10.7 |
| Kinesin$^{EL-CC-Di-IR}$ | 69.7 | 21.5 |
| Kinesin$^{EL-A-Ala}$ | 40 | 14 |
| Kinesin$^{EL-A-Ala\ +\ Peptide\ B}$ | 69 | 19.1 |

| **Table S3. SEC-SAXS Data collection parameters** | |
|---|---|
| **Source Type** | Bending magnet |
| **Beamline** | B21, biological X-ray scattering |
| **Detector** | Eiger 4M (Dectris) |
| **Beam size at the focal point at detector (FWHM)** | 34 x 40 µm (horizontal vertical) |
| **Beam size at sample** | 01102 x 240 µm |
| **Wavelength range** | 0.89 – 1.3 Å |
| **Mode/Column** | SEC/Superose 6 (10/300) |
| **Exposure time** | 3 s |

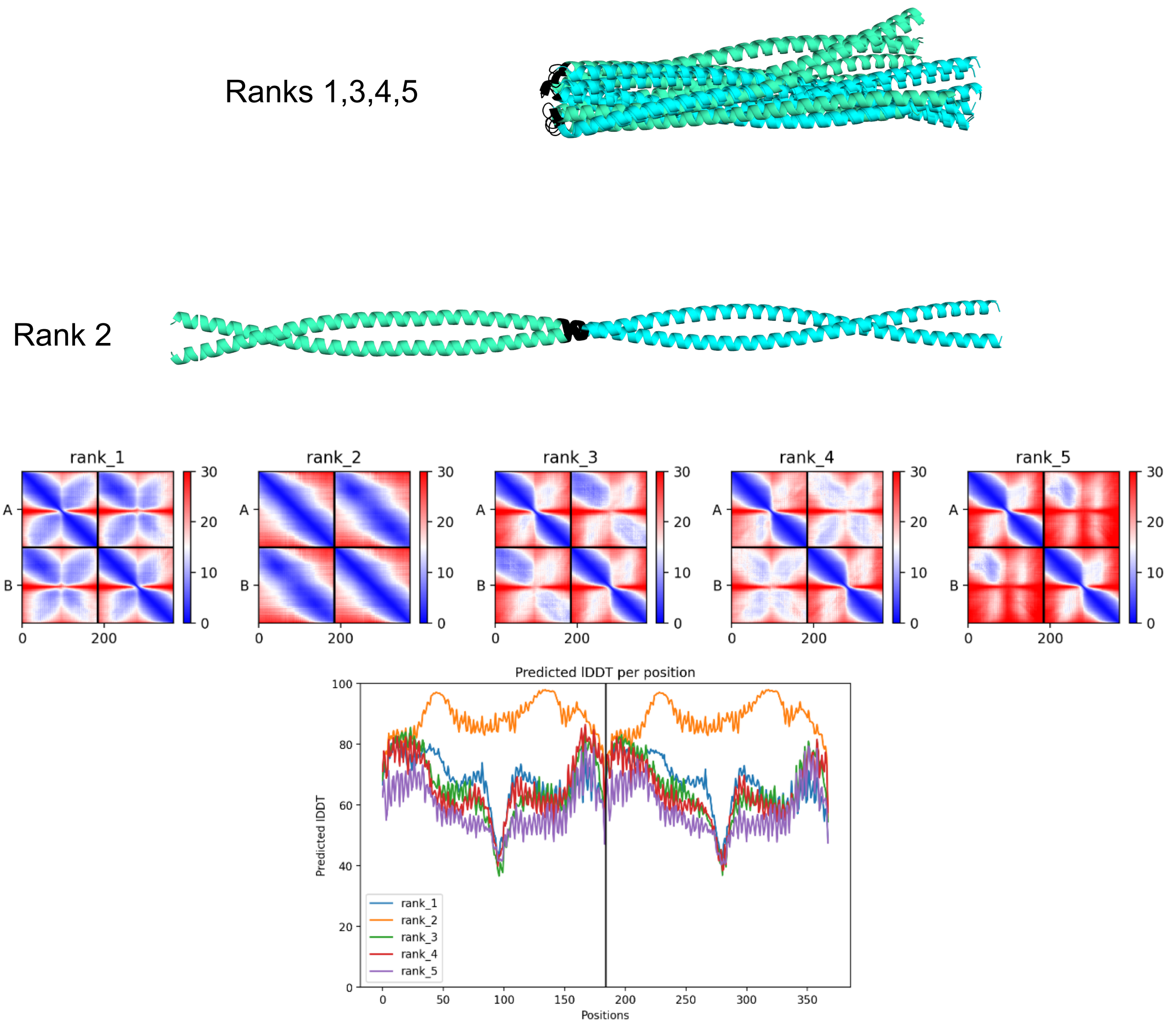


Sequence:
`iskmksevkslvnrskqlesaqtdsnrkmnaserelaacqllisqheakiksltdymqnmeqkrrqle`
`esqdslseelaklraqekmhevsfqdkekehltrlqdaeevkkaleqqmeshreahqkqlsrlrdeie`
`ekqriideirdlnqklqleqerlssdynklkiedqerevklekllllln`

**Fig. S1. Alignment of AlphaFold2 models of Kinesin-1$^{WT}$ CC2-elbow-CC3 with corresponding PAE and pLDDT plots and sequence**. 4 aligned models (ranks 1,3,4,5) of Kinesin-1$^{WT}$ CC2-elbow-CC3 have a folded structural prediction with a flexible elbow (black) between CC2 (teal) and CC3 (cyan) The remaining model predicts an extended coiled coil conformation (rank 2).

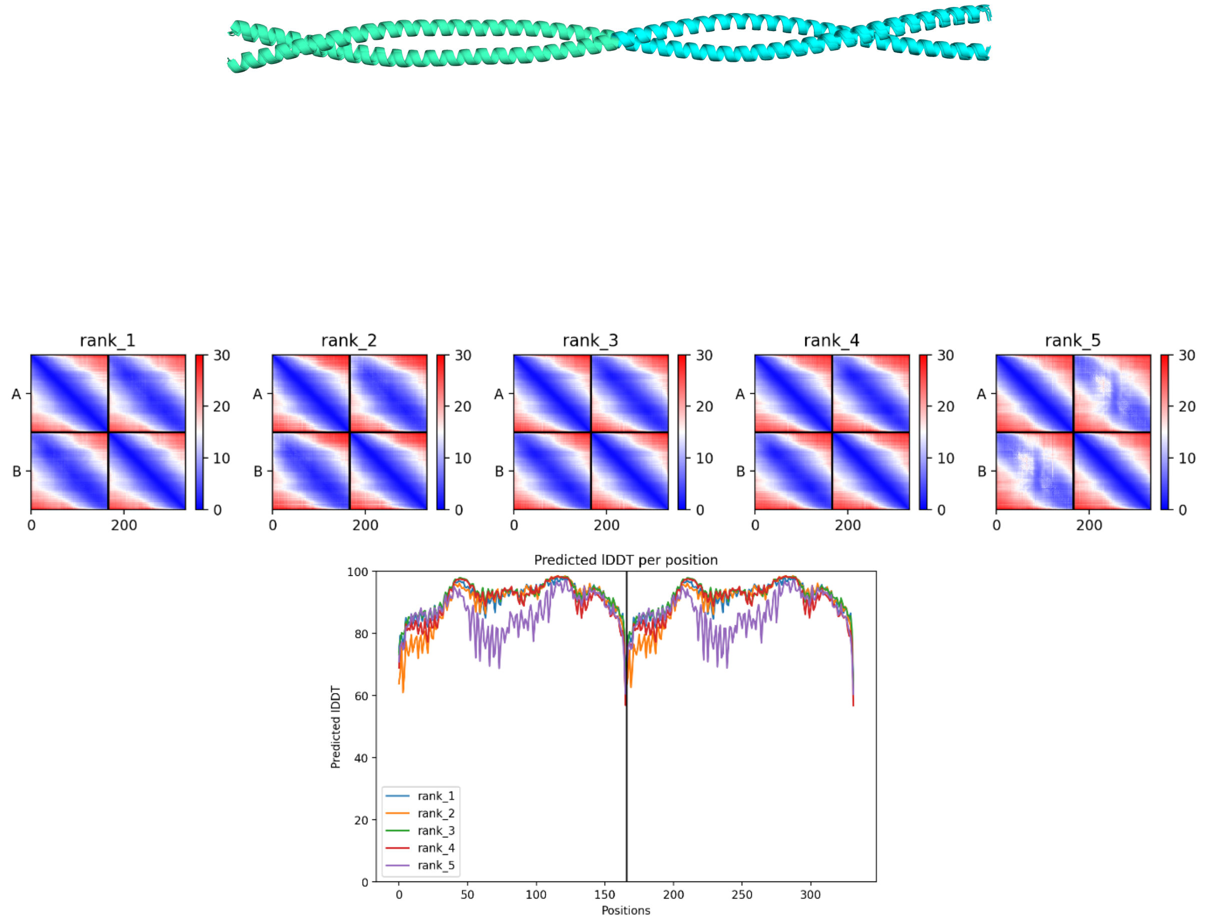


Sequence:
iskmksevkslvnrskqlesaqtdsnrkmnaserelaacqllisqheakiksltdymqnmeqkrrqle
esqdslseelaklraqlqdaeevkkaleqqmeshreahqkqlsrlrdeieekqriideirdlnqklql
eqerlssdynklkiedqerevkleklllln

**Fig. S2. Alignment of AlphaFold2 models of Kinesin-1$^{\text{Delta Elbow}}$ CC2-elbow-CC3 with corresponding PAE and pLDDT plots and sequence.** All five models closely overlay with the same extended coiled coil structural prediction, CC2 (teal) and CC3 (cyan).

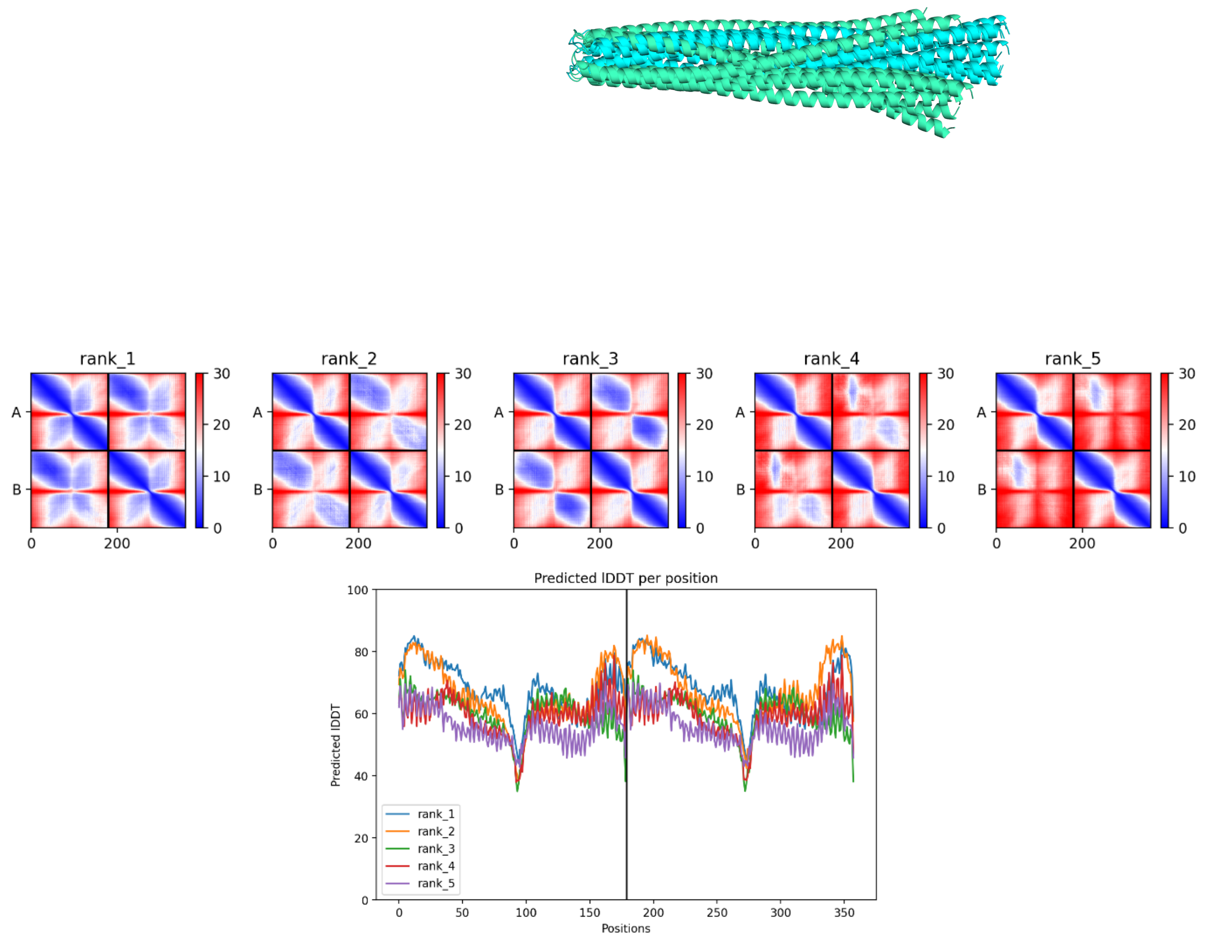


Sequence:

`iskmksevkslvnrskqlesaqtdsnrkmnaserelaacqllisqheakiksltdymqnmeqkrrqle`
`esqdslseelaklraqekmhevsfqdktrlqdaeevkkaleqqmeshreahqkqlsrlrdeieekqri`
`ideirdlnqklqleqerlssdynklkiedqerevklekllllln`

**Fig. S3. Alignment of AlphaFold2 models of Kinesin-1$^{\mathrm{Elbow\ Lock}}$ CC2-elbow-CC3 with corresponding PAE and pLDDT plots and sequence.** All five models have a folded structural prediction with a flexible hinge between CC2 (teal) and CC3 (cyan).

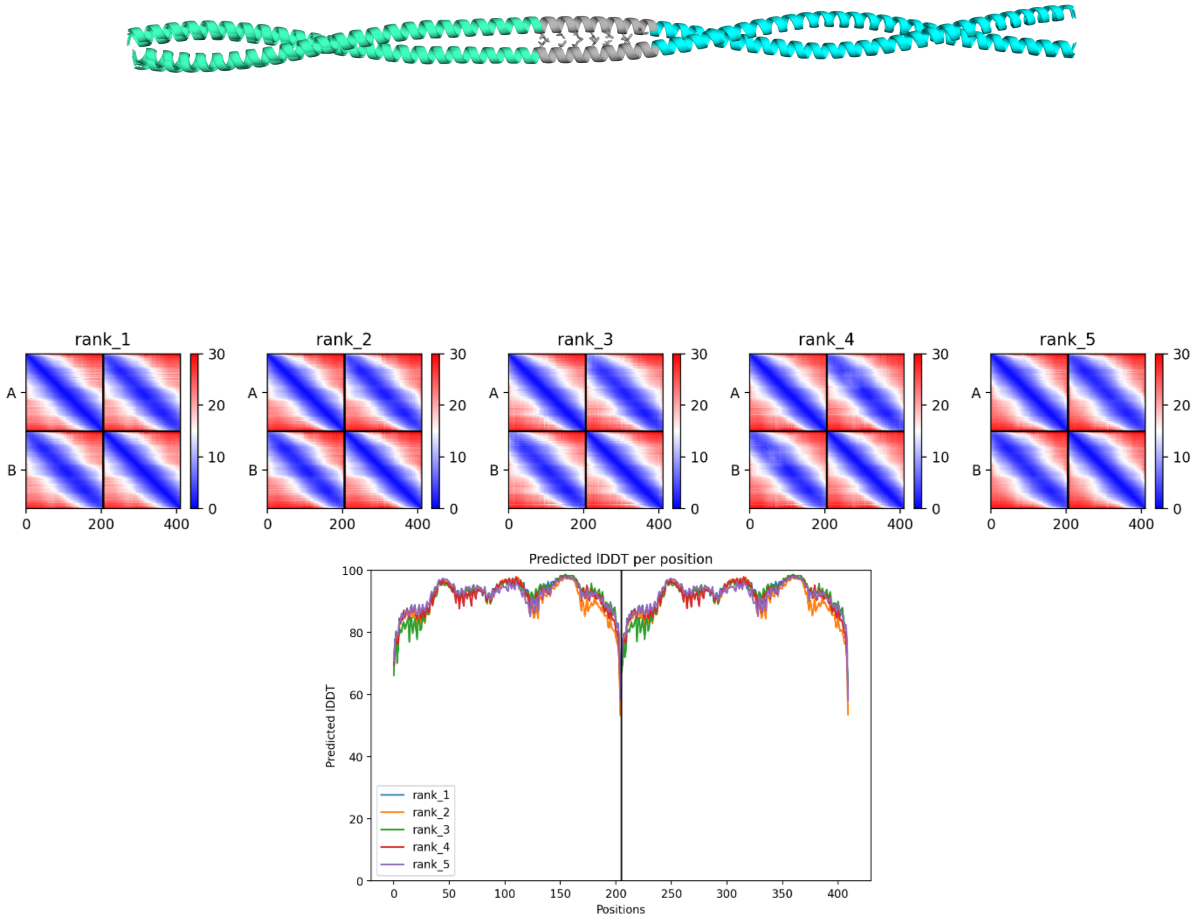


Sequence:
iskmksevkslvnrskqlesaqtdsnrkmnaserelaacqllisqheakiksltdymqnmeqkrrqle
esqdslseelaklraqekmhevsfqdkKQEIAAIKKENAALKWEIAALKQEIAtrlqdaeevkkaleq
qmeshreahqkqlsrlrdeieekqriideirdlnqklqleqerlssdynklkiedqerevklekllll
n

**Fig. S4. Alignment of AlphaFold2 models of Kinesin-1$^{EL\text{-}CC\text{-}Di\text{-}IR}$ CC2-elbow-CC3 with corresponding PAE and pLDDT plots and sequence.** All five models closely overlay with the same extended coiled coil structural prediction, CC2 (teal) and CC3 (cyan) with CC-Di in grey. Side chains are shown as sticks for the hydrophobic core of CC-Di.

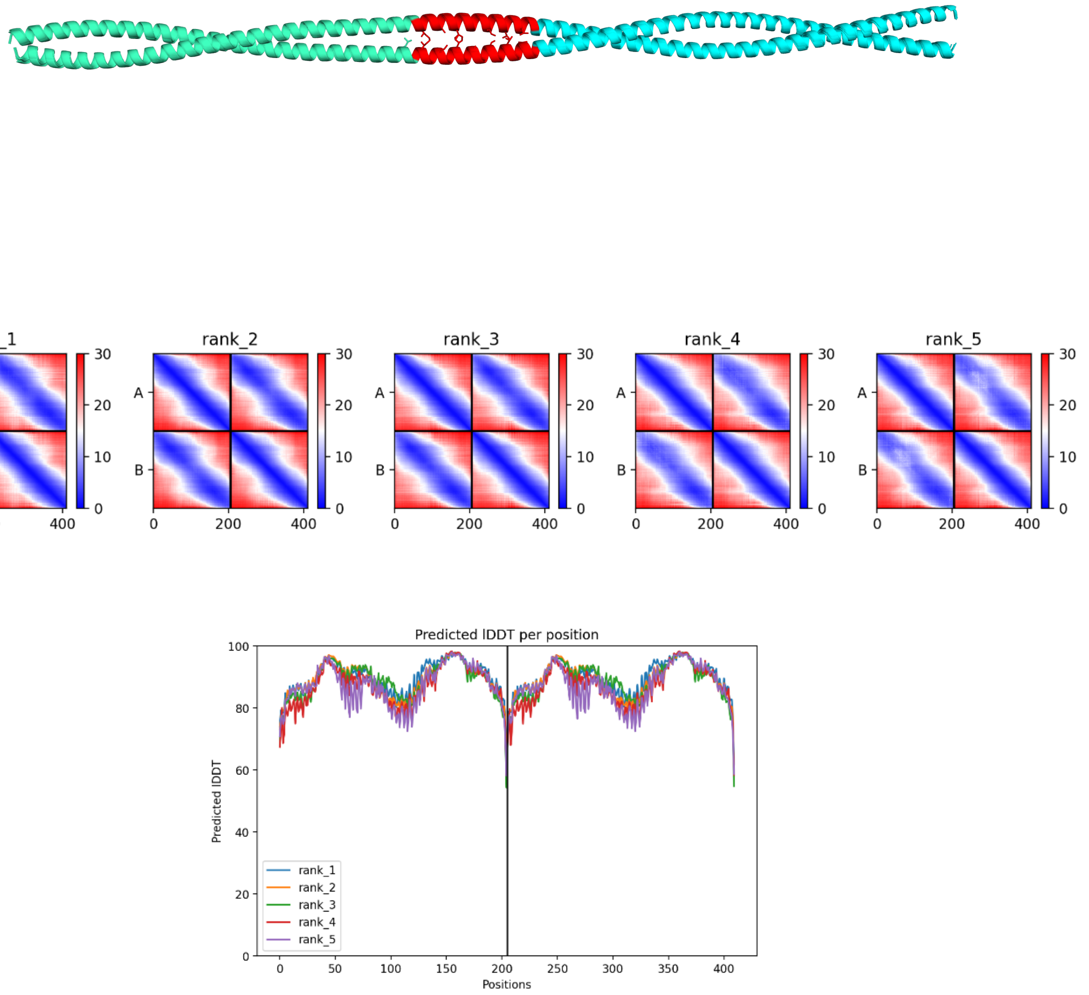


Sequence:

iskmksevkslvnrskqlesaqtdsnrkmnaserelaacqllisqheakiksltdymqnmeqkrrqle
esqdslseelaklraqekmhevsfqdkEQELAALDQEIAAAEQELAALDWQIQtrlqdaeevkkaleq
qmeshreahqkqlsrlrdeieekqriideirdlnqklqleqerlssdynklkiedqerevklekllll
n

**Fig. S5. Alignment of AlphaFold2 models of Kinesin-1$^{EL\text{-}pA\text{-}IR}$ CC2-elbow-CC3 with corresponding PAE and pLDDT plots and sequence.** All five models closely overlay with the same extended coiled coil structural prediction, CC2 (teal) and CC3 (cyan) with pA in red. Side chains are shown as sticks for the hydrophobic core of pA.

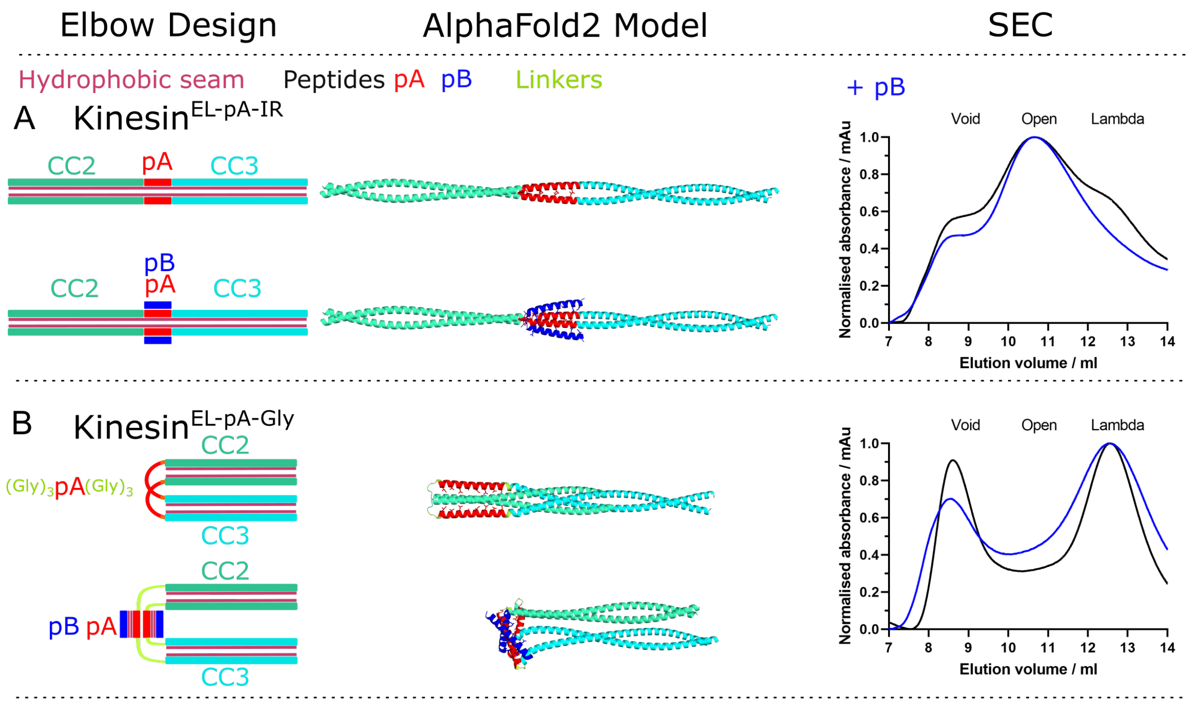


**Fig. S6. Designing an allosteric switch into kinesin-1 that is activated by a *de novo* designed peptide.** From left to right: cartoon illustrations of each elbow design without (top) and with (bottom) peptide pB; AlphaFold2 models of KHC for the CC2-linker-CC3 region without (top) and with (bottom) peptide pB; and size-exclusion chromatography (SEC) elution profiles of purified heterotetrameric KHC-KLC complexes without (black) and with (blue) peptide pB. Colour scheme for the structural cartoons: KHC CC2 teal, CC3 cyan; hydrophobic seams/cores, pink; peptide pA, red; linkers, yellow; and peptide pB, blue. **A.** Insertion of pA in register (IR) between CC2 and CC3 drives helical read through and favours the open state without or with pB. **B.** Insertion of pA between flexible $(Gly)_3$ linkers after CC2 and before CC3 favours the lambda particle without or with pB.

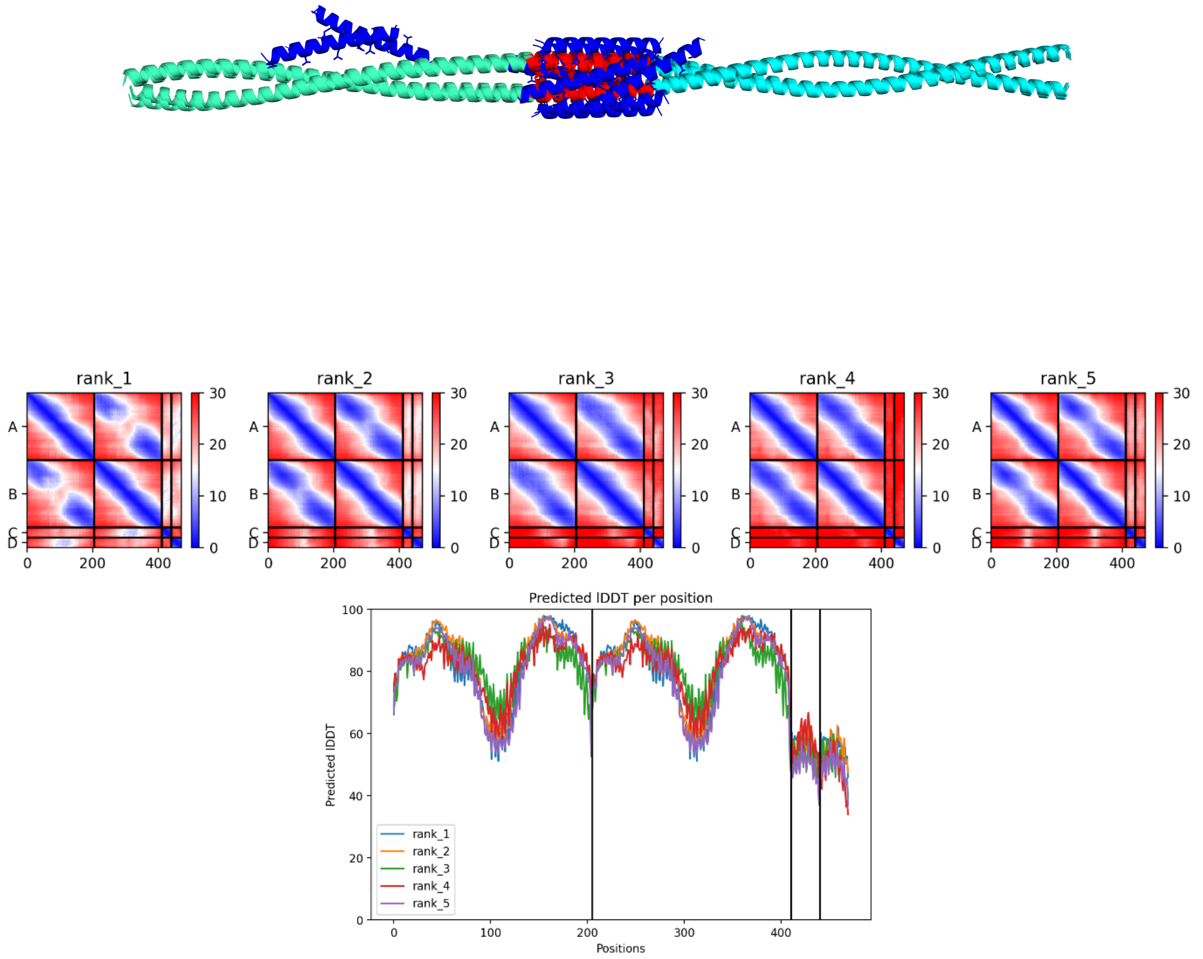


Sequence:

iskmksevkslvnrskqlesaqtdsnrkmnaserelaacqllisqheakiksltdymqnmeqkrrqle
esqdslseelaklraqekmhevsfqdkEQELAALDQEIAAAEQELAALDWQIQtrlqdaeevkkaleq
qmeshreahqkqlsrlrdeieekqriideirdlnqklqleqerlssdynklkiedqerevklekllll
n

Peptide pB: GQLKQRRAALKQRIAALKQRRAALKWQIQG

**Fig. S7. Alignment of AlphaFold2 models of Kinesin-1$^{EL\text{-}pA\text{-}IR}$ CC2-elbow-CC3 with peptide pB with corresponding PAE and pLDDT plots and sequences.** All five models of Kinesin-1$^{EL\text{-}pA\text{-}IR}$ CC2-elbow-CC3 closely overlay with the same extended coiled coil structural prediction, CC2 (teal) and CC3 (cyan) with pA in red. The placement of peptide pB (blue) is more varied. Side chains are shown as sticks for the hydrophobic core of pA and pB.

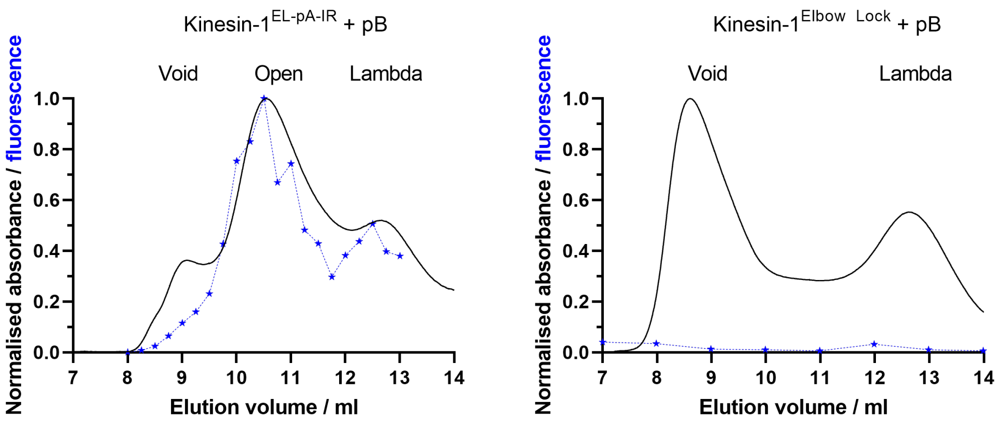


**Fig. S8. Binding of peptide pB to Kinesin-1$^{EL\text{-}pA\text{-}IR}$ is observed by fluorescence in SEC (left) compared to no binding to Kinesin-1$^{Elbow\ Lock}$ (right).** Peptide pB was added to purified Kinesin-1 (Kif5C, KLC1) protein at 4ºC for 1h with agitation and then subject to SEC. Elution of protein in SEC was monitored by absorbance at 280 nm and normalised (black curve). Presence of peptide-B was monitored by fluorescence of TAMRA at 555 nm in individual fractions using a plate reader and normalised (blue stars). Coelution of peptide pB and Kinesin-1$^{EL\text{-}pA\text{-}IR}$ indicates binding.

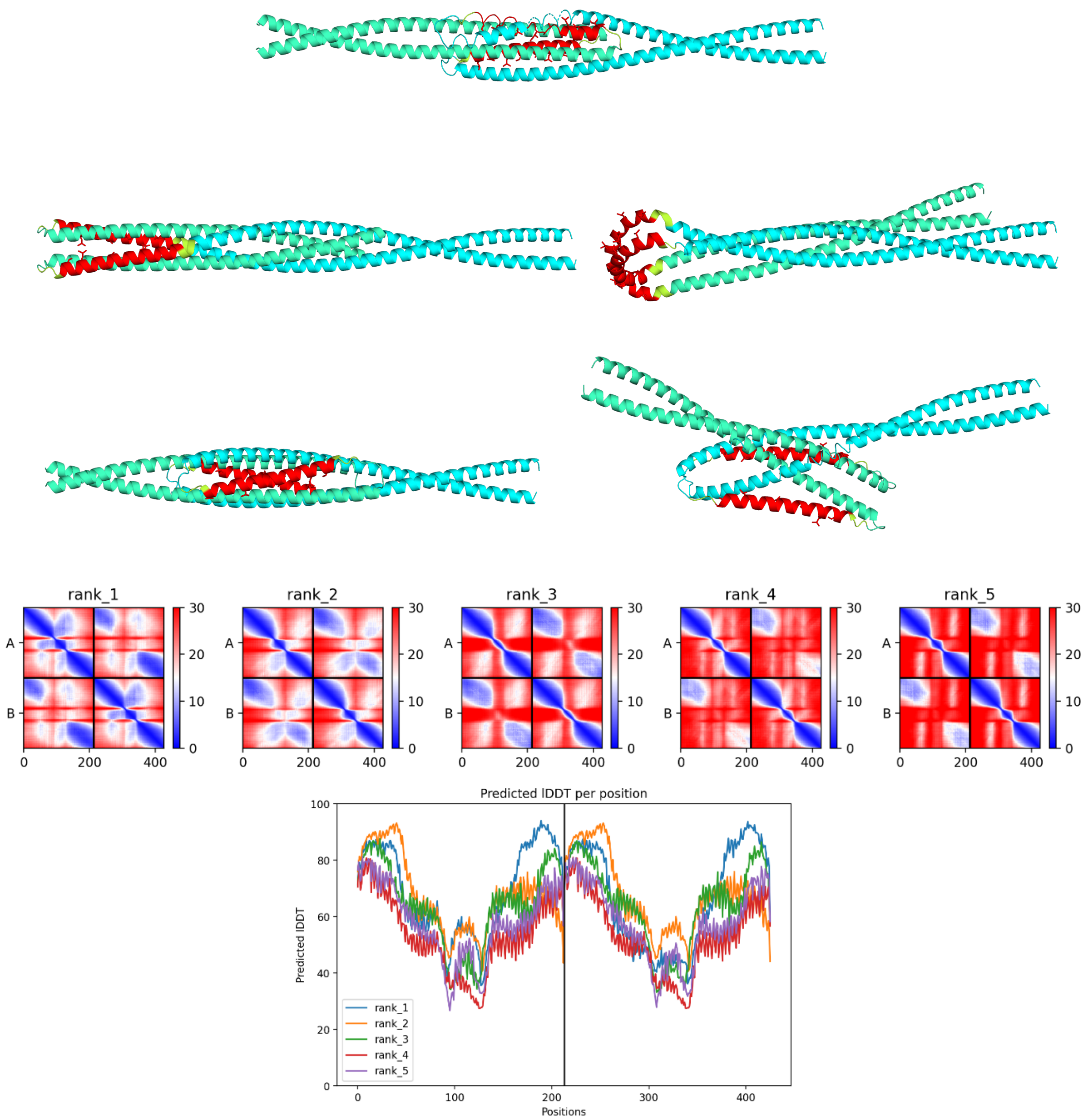


Sequence:

```
iskmksevkslvnrskqlesaqtdsnrkmnaserelaacqllisqheakiksltdymqnmeqkrrqle
esqdslseelaklraqekmhevsfqdkGGGQLEQELAALDQEIAAAEQELAALDWQIQGGGtrlqdae
evkkaleqqmeshreahqkqlsrlrdeieekqriideirdlnqklqleqerlssdynklkiedqerev
kleklllln
```

**Fig. S9. AlphaFold2 models of Kinesin-1$^{EL\text{-}pA\text{-}Gly}$ CC2-elbow-CC3 with corresponding PAE and pLDDT plots and sequence.** The five models vary but all have a folded structural prediction with a flexible hinge between CC2 (teal) and CC3 (cyan) with pA in red, and GGG linkers in yellow. Side chains are shown as sticks for the hydrophobic core of pA.

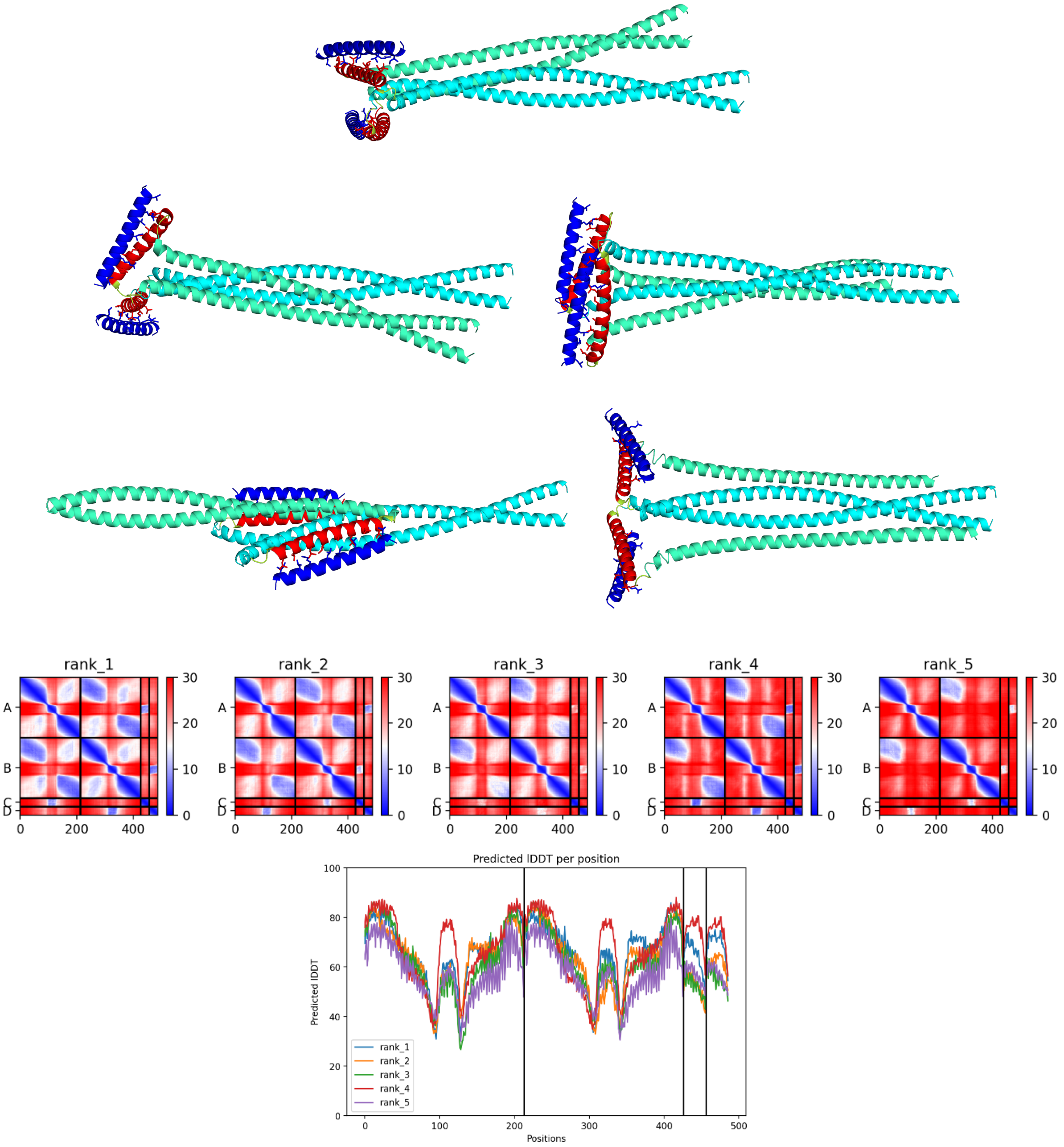


Sequence:

```
iskmksevkslvnrskqlesaqtdsnrkmnaserelaacqllisqheakiksltdymqnmeqkrrqle
esqdslseelaklraqekmhevsfqdkGGGQLEQELAALDQEIAAAEQELAALDWQIQGGGtrlqdae
evkkaleqqmeshreahqkqlsrlrdeieekqriideirdlnqklqleqerlssdynklkiedqerev
klekllln
```

Peptide pB: GQLKQRRAALKQRIAALKQRRAALKWQIQG

**Fig. S10. AlphaFold2 models of Kinesin-1$^{EL\text{-}pA\text{-}Gly}$ CC2-elbow-CC3 with peptide B with corresponding PAE and pLDDT plots and sequences** The five models vary but all have a folded structural prediction with a flexible hinge between CC2 (teal) and CC3 (cyan) accommodating pA (red) binding to pB (blue). GGG linkers are in yellow and side chains are shown as sticks for the hydrophobic core of pA and pB.

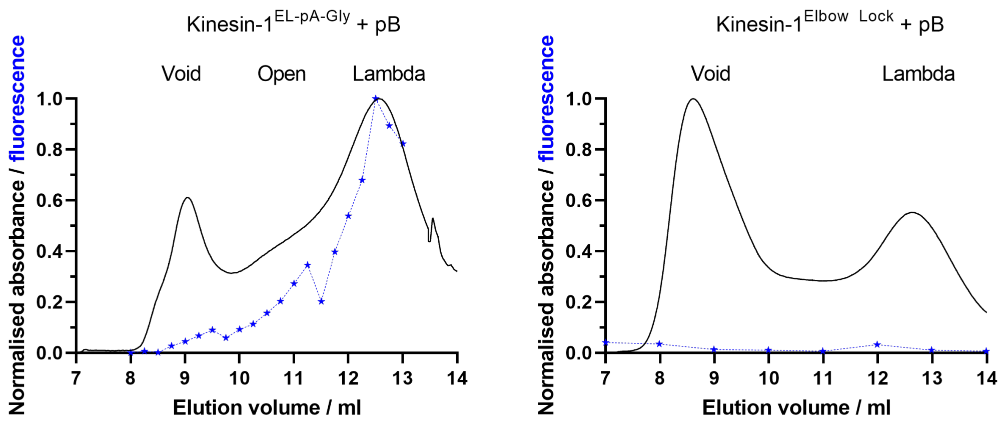


**Fig. S11. Binding of peptide pB to Kinesin-1$^{EL\text{-}pA\text{-}Gly}$ is observed by fluorescence in SEC (left) compared to no binding to Kinesin-1$^{Elbow\ Lock}$ (right).** Peptide pB was added to purified Kinesin-1 (Kif5C, KLC1) protein at 4°C for 1h with agitation and then subject to SEC. Elution of protein in SEC was monitored by absorbance at 280 nm and normalised (black curve). Presence of peptide-B was monitored by fluorescence of TAMRA at 555 nm in individual fractions using a plate reader and normalised (blue stars). Coelution of peptide pB and Kinesin-1$^{EL\text{-}pA\text{-}Gly}$ indicates binding.

Ranks 1-4

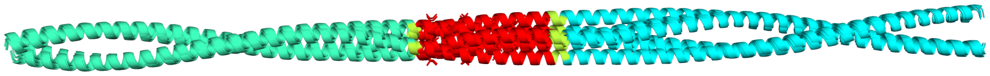

Ranks 5

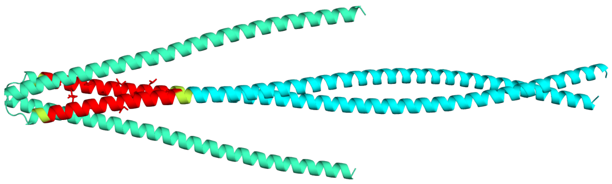

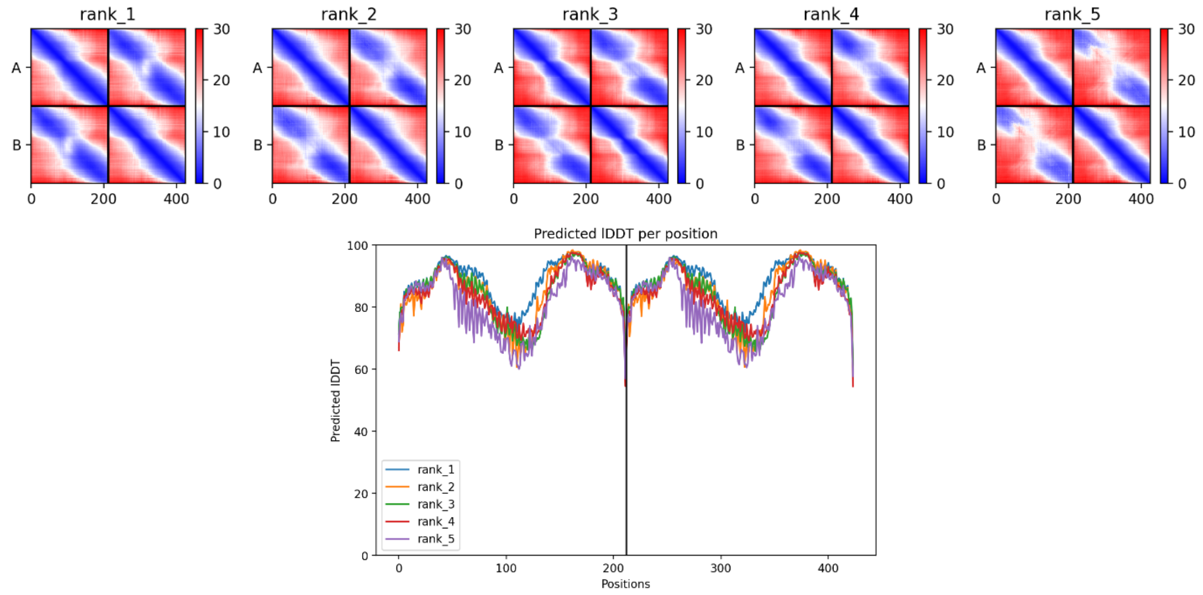


Sequence:

Iskmksevkslvnrskqlesaqtdsnrkmnaserelaacqllisqheakiksltdymqnmeqkrrqle
esqdslseelaklraqekmhevsfqdkAAAQLEQELAALDQEIAAAEQELAALDWQIQAAtrlqdaee
vkkaleqqmeshreahqkqlsrlrdeieekqriideirdlnqklqleqerlssdynklkiedqerevk
lekllllln

**Fig. S12. Alignment of AlphaFold2 models of Kinesin-1$^{EL\text{-}pA\text{-}Ala}$ CC2-elbow-CC3 with corresponding PAE and pLDDT plots and sequence.** 4 aligned models (ranks 1-4) of Kinesin-1$^{EL\text{-}pA\text{-}Ala}$ CC2-elbow-CC3 closely overlay with the same extended coiled coil structural prediction of CC2 (teal) and CC3 (cyan) with peptide-A in red and AAA/AA linkers in yellow. The final model (rank 5) predicts a folded conformation. Side chains are shown as sticks for the hydrophobic core of pA.

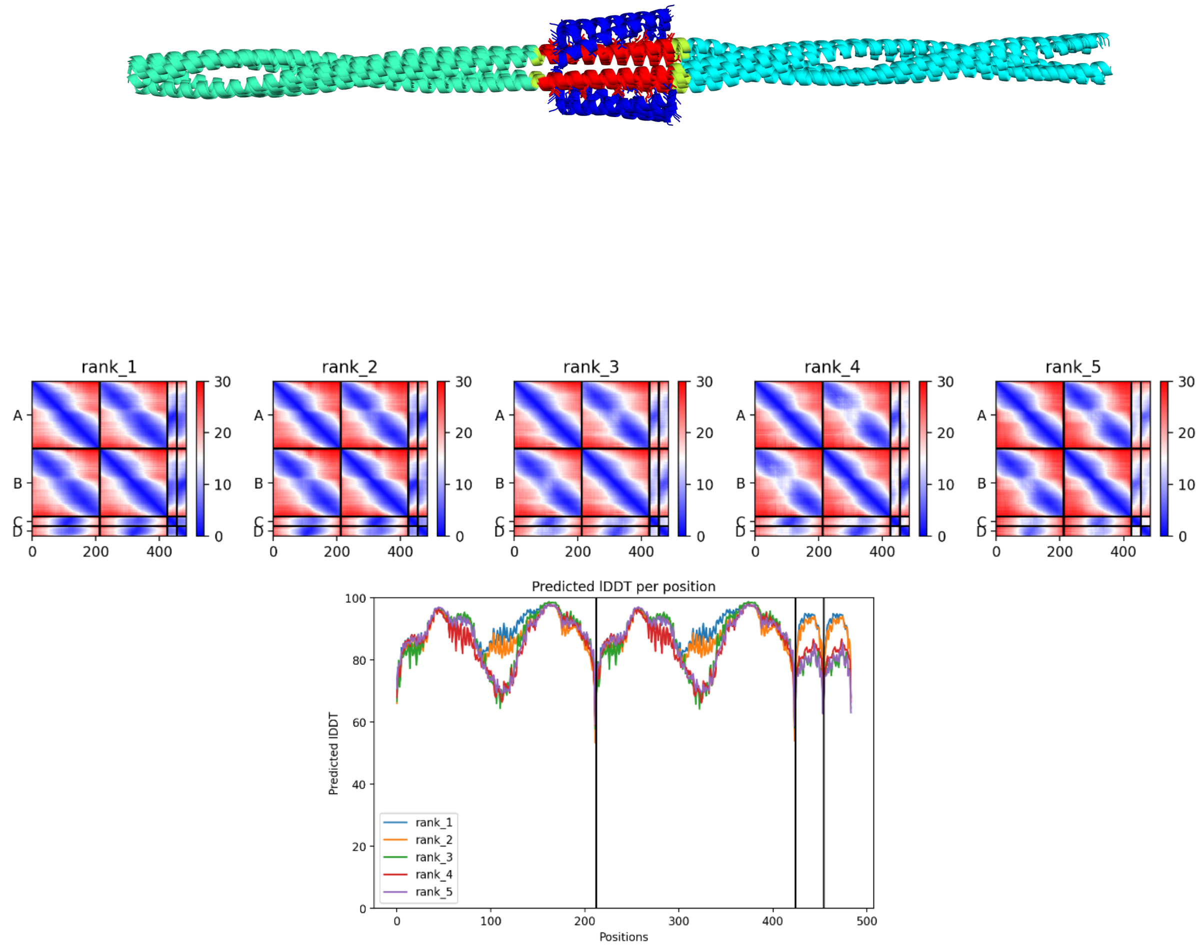


Sequence:

Iskmksevkslvnrskqlesaqtdsnrkmnaserelaacqllisqheakiksltdymqnmeqkrrqle
esqdslseelaklraqekmhevsfqdkAAAQLEQELAALDQEIAAAEQELAALDWQIQAAtrlqdaee
vkkaleqqmeshreahqkqlsrlrdeieekqriideirdlnqklqleqerlssdynklkiedqerevk
leklllln

Peptide pB: GQLKQRRAALKQRIAALKQRRAALKWQIQG

**Fig. S13. Alignment of AlphaFold2 models of Kinesin-1$^{EL-pA-Ala}$ CC2-elbow-CC3 with peptide B with corresponding PAE and pLDDT plots and sequences.** All five models of Kinesin-1$^{EL-pA-IR}$ CC2-elbow-CC3 and peptide pB closely overlay with the same extended coiled coil structural prediction of CC2 (teal) and CC3 (cyan) with pA in red, peptide pB in blue and AAA/AA linkers in yellow. Side chains are shown as sticks for the hydrophobic core of pA and pB.

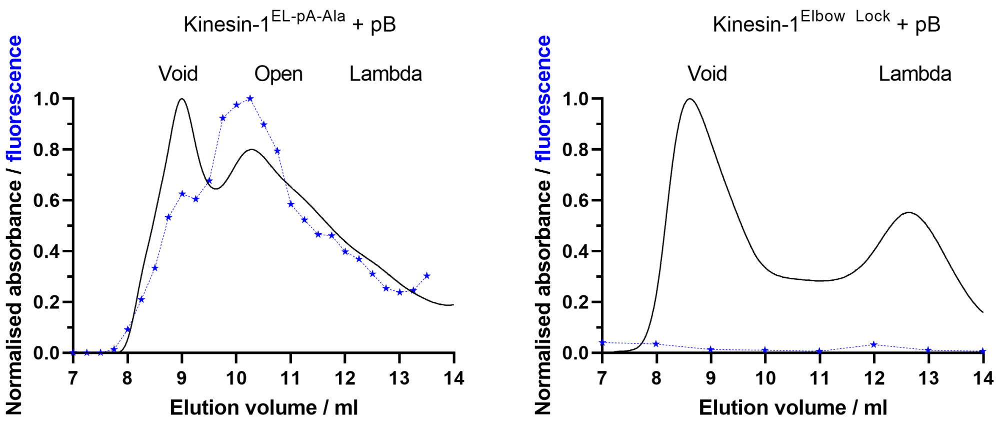


**Fig. S14. Binding of peptide pB to Kinesin-1$^{EL\text{-}pA\text{-}Ala}$ is observed by fluorescence in SEC (left) compared to no binding to Kinesin-1$^{Elbow\ Lock}$ (right).** Peptide-B was added to purified Kinesin-1 (Kif5C, KLC1) protein at 4°C for 1h with agitation and then subject to SEC. Elution of protein in SEC was monitored by absorbance at 280 nm and normalised (black curve). Presence of peptide pB was monitored by fluorescence of TAMRA at 555 nm in individual fractions using a plate reader and normalised (blue stars). Coelution of peptide pB and Kinesin-1$^{EL\text{-}pA\text{-}Ala}$ indicates binding.

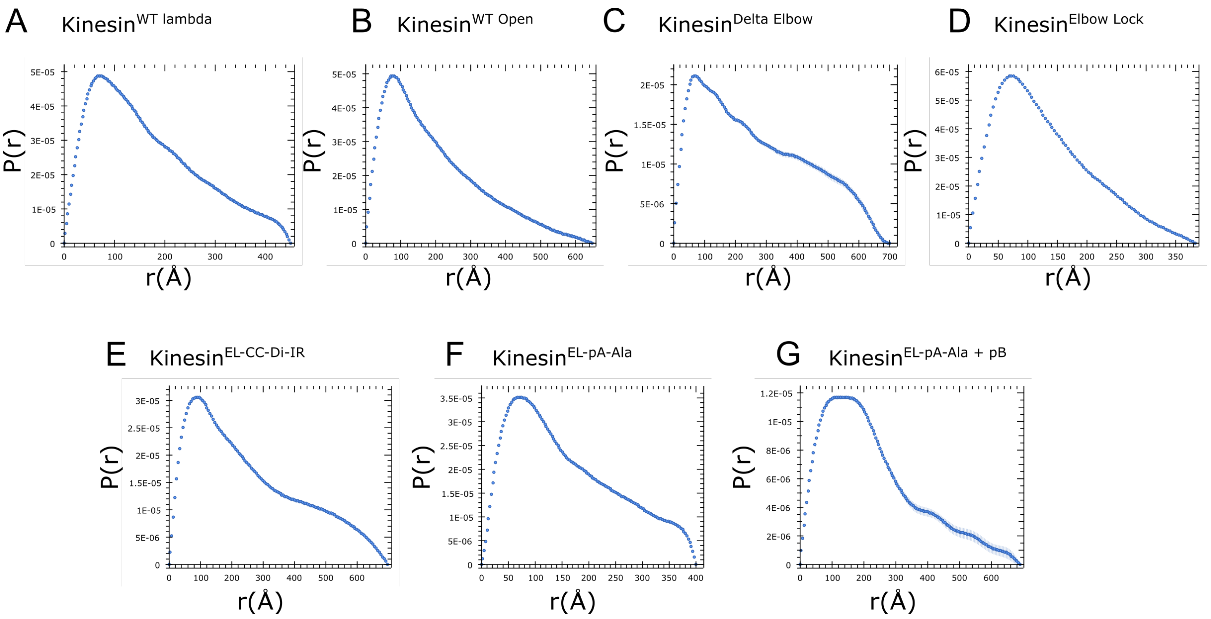


**Fig. S15. Interatomic distance distribution plots for kinesin-1 variants from SEC-SAXS scattering data. A.** Kinesin-1$^{WT\ Lambda}$, **B.** Kinesin-1$^{WT\ Open}$ **C.** Kinesin-1$^{Delta\ Elbow}$, **D.** Kinesin-1$^{Elbow\ Lock}$, **E.** Kinesin-1$^{EL\text{-}CC\text{-}Di\text{-}}$, **F.** Kinesin-1$^{EL\text{-}pA\text{-}Ala}$, **G.** Kinesin-1$^{EL\text{-}pA\text{-}Ala}$ + pB.

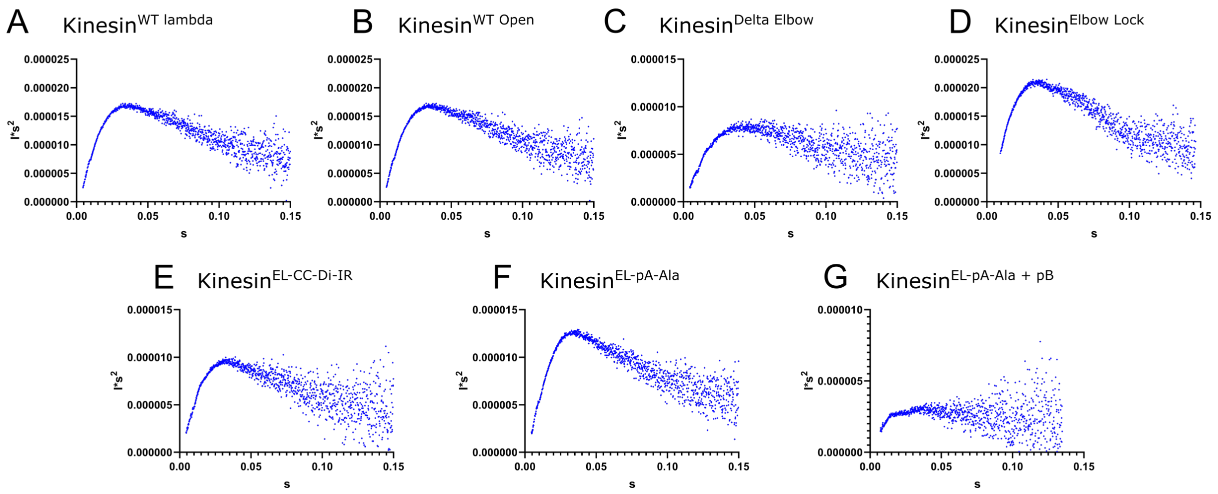


**Fig. S16. Kratky plot indicated flexibility in all the kinesin-1 complexes from SEC-SAXS scattering data.** Kratky plots give a qualitative assessment of globularity or flexibility in samples. Whilst globular proteins are expected to converge with a gaussian peak, these plots show a plateau at high s indicating flexibility in the complex.

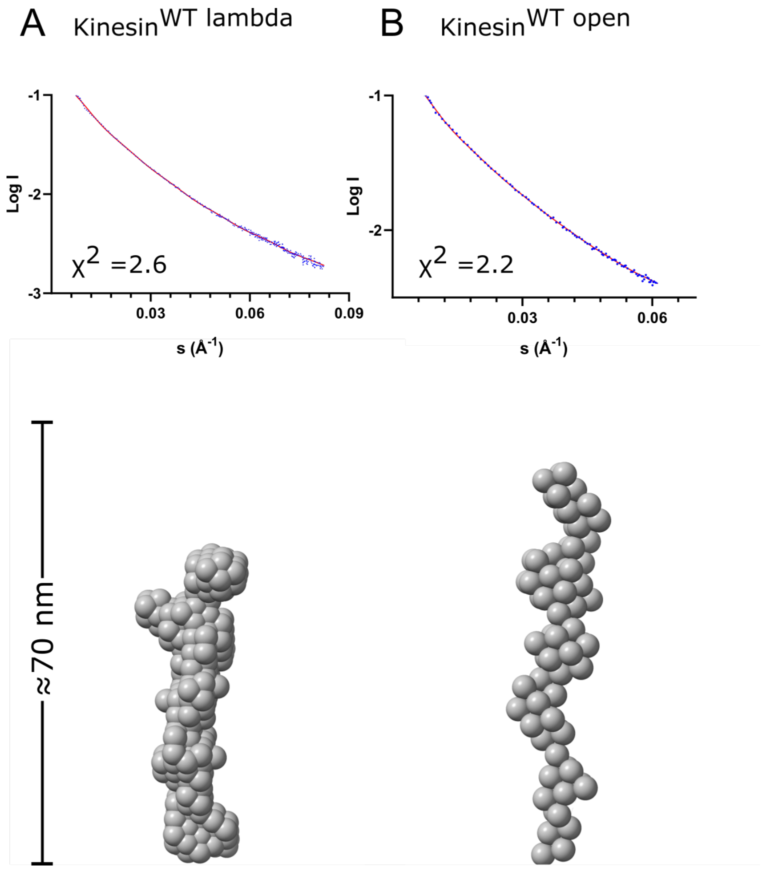


**Fig. S17. SEC-SAXS in-solution shape determination and conformational states of Kinesin-1$^{WT}$.** Top: Comparison of trimmed experimental scattering profiles (blue) with theoretical scattering profiles (red) generated from ab-initio models using DAMMIN. The quality of fit is assessed using the $\chi^2$ statistic. Bottom: Representative models obtained from DAMMIN illustrating the shapes of the kinesin-1 variants, **A.** Kinesin-1$^{WT\ lambda}$ is in the lambda state $D_{max}$ = 45 nm $\chi^2$=2.6, **B.** Kinesin-1$^{WT\ open}$ is in the open state $D_{max}$ = 64.7 nm $\chi^2$ = 2.2.
